# Phylogenetic reconstruction of myeloproliferative neoplasm reveals very early origins and lifelong evolution

**DOI:** 10.1101/2020.11.09.374710

**Authors:** Nicholas Williams, Joe Lee, Luiza Moore, E Joanna Baxter, James Hewinson, Kevin J Dawson, Andrew Menzies, Anna L Godfrey, Anthony R Green, Peter J Campbell, Jyoti Nangalia

**Author notes:** senior authors (Anthony R Green, Peter J Campbell, Jyoti Nangalia). Corresponding author: Jyoti Nangalia, Wellcome Sanger Institute, Hinxton, UK.

## Abstract

Mutations in cancer-associated genes drive tumour outgrowth. However, the timing of driver mutations and dynamics of clonal expansion that lead to human cancers are largely unknown. We used 448,553 somatic mutations from whole-genome sequencing of 843 clonal haematopoietic colonies to reconstruct the phylogeny of haematopoiesis, from embryogenesis to clinical disease, in 10 patients with myeloproliferative neoplasms which are blood cancers more common in older age. *JAK2*^*V617F*^, the pathognomonic mutation in these cancers, was acquired *in utero* or childhood, with upper estimates of age of acquisition ranging between 4.1 months and 11.4 years across 5 patients. *DNMT3A* mutations, which are associated with age-related clonal haematopoiesis, were also acquired *in utero* or childhood, by 7.9 weeks of gestation to 7.8 years across 4 patients. Subsequent driver mutation acquisition was separated by decades. The mean latency between *JAK2*^*V617F*^ acquisition and clinical presentation was 31 years (range 12-54 years). Rates of clonal expansion varied substantially (<10% to >200% expansion/year), were affected by additional driver mutations, and predicted latency to clinical presentation. Driver mutations and rates of expansion would have been detectable in blood one to four decades before clinical presentation. This study reveals how driver mutation acquisition very early in life with life-long growth and evolution drive adult blood cancer, providing opportunities for early detection and intervention, and a new paradigm for cancer development.

## INTRODUCTION

Human cancers harbor hundreds to hundreds of thousands of somatically acquired DNA mutations. Whilst the majority of such mutations do not affect the cancer’s biology, a minority drive tumour initiation, growth and progression^1^. These so-called driver mutations occur in recurrently mutated cancer genes, and stimulate the cell acquiring it to expand into a clone. With a large enough clonal expansion, typically abetted by acquisition of further driver mutations, a cancer emerges. Little is known about what ages driver mutations occur, the timelines of clonal expansion over a patient’s lifetime, or how these relate to clinical presentation with overt cancer. Some mutational processes accrue at a constant rate across life, representing a ‘molecular clock’^2,3^. Knowing this tissue-specific rate of mutation accumulation, it has been possible to infer broad estimates for the timing of driver mutations for some cancers^4,5^.

In patients with blood cancers, the observation of normal blood counts months to years prior to diagnosis has led to the prediction that tumour development occurs quickly, and therefore driver mutations must occur late in life. Estimates from cancer incidences in Japanese atomic survivors who developed chronic myeloid leukaemia have suggested a mean latency time of only 8 years between *BCR-ABL1* induction and clinical presentation^6^. However, the presence of driver mutations in normal tissues^7–12^, including blood from healthy individuals who harbor age-related clonal haematopoiesis (CH)^13–17^, some of whom subsequently develop malignancies, supports a longer multi-hit evolutionary trajectory of cancer. Understanding the absolute timelines of cancer evolution is critical for efforts aimed at early detection and intervention, especially if a given cancer takes decades to emerge after its first driver mutation.

Myeloproliferative neoplasms (MPN) are blood cancers driven by somatic driver mutations in haematopoietic stem cells (HSC) that result in increased mature myeloid cell production^18^. Most patients harbor *JAK2*^*V617F*^, and this can either be the only driver mutation or occur in co-operation with driver mutations in other genes such as *DNMT3A* or *TET2*^19^. The clinical course often spans decades, with phenotypic disease progression occurring upon acquisition of additional driver mutations^20^. MPNs provide a unique opportunity to capture the earliest stages of tumourigenesis through to disease evolution which are otherwise inaccessible in other malignancies. Here, we undertake whole-genome sequencing (WGS) of individual single-cell derived haematopoietic colonies, and targeted resequencing of longitudinal blood samples from patients with MPN, to assess the absolute timing of driver mutations, tumour evolutionary dynamics, and the fundamental nature of driver mutation mediated clonal selection *in vivo*.

## RESULTS

### Using somatic mutations for haematopoietic lineage tracing in patients with MPN

The mutations present in a somatic cell’s genome have accumulated throughout its ancestral lineage, passed from mother to both daughter cells with each cell division. We identified somatic mutations in individual HSCs from patients with MPN, using them to reconstruct the lineage relationships among both malignant and normal blood cells in each patient^21^. Given that somatic mutation burden does not differ between HSCs and myeloid progenitors^21,22^, we undertook WGS of *in-vitro* expanded single-cell derived haematopoietic colonies as faithful surrogates for the genomes of their parental HSCs. We then ‘recaptured’ these somatic mutations in bulk peripheral blood cells using targeted sequencing in order to longitudinally track clones and infer population estimates (Fig.1a, b).

**Figure 1.**
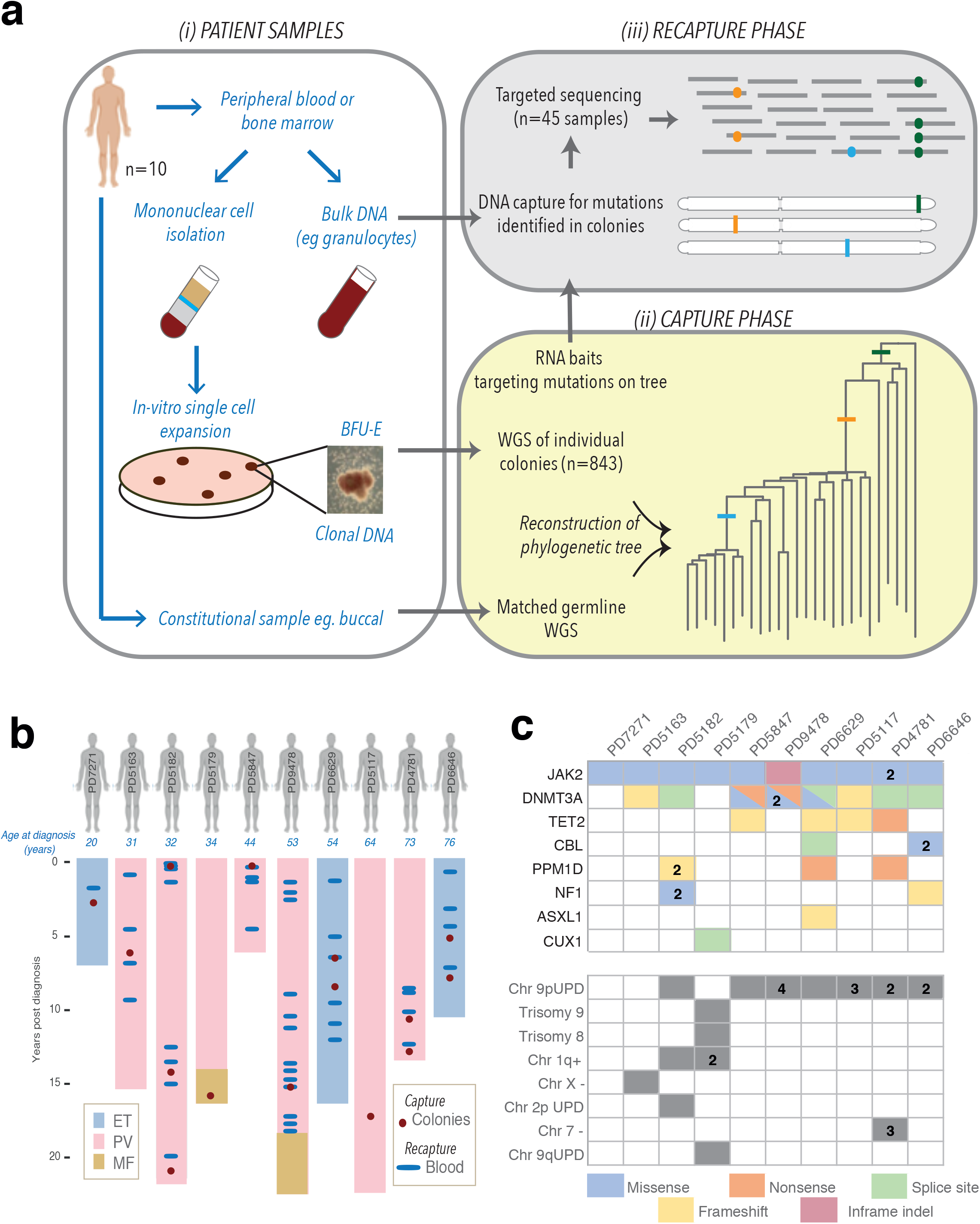
Patient cohort and experimental design. A. Experimental design. WGS, whole genome sequencing; BFU-E, Burst forming unit-erythroid. B. Patient cohort showing ages at diagnosis, disease phase and duration of disease, sample types and timepoints. ET, Essential thrombocythemia; PV polycythemia vera; MF, myelofibrosis. The length of the shaded bars represents the duration of disease, either to last follow-up or to patient death. C. Driver mutations, both single nucleotide variants and insertions/deletions, as well as copy number aberrations identified in at least one colony within each patient are shown. Shaded colours represent the type of mutation and the numbers within the squares represent the number of mutations or copy number aberrations in individual patients.

Our cohort comprised 10 patients with MPN diagnosed between ages 20 and 76 years (3 essential thrombocythemia (ET), 5 polycythemia vera (PV) and 2 post-PV myelofibrosis (MF)) (Fig.1b, Extended Table 1). We obtained 952 colonies from 15 timepoints spanning both diagnosis and disease course for WGS to a mean depth of ~14x (Fig.1b). Colonies with low sequencing coverage or evidence of non-clonality were excluded, and 843 were included in the final analyses (Extended Table 2). We identified 448,553 somatic single nucleotide variants (SNV) and 14,851 small insertions and deletions. Variant allele fractions (VAFs) clustered around 0.5 (Extended Fig.1a), confirming colonies derived from a single cell. There were no additional subclonal peaks in the VAF distribution confirming that few mutations were acquired *in-vitro* (Extended Fig.1b). All patients harbored mutated-*JAK2* (9 *JAK2*^*V617F*^, 1 *JAK2*^exon^ ^12^), and 9 patients had additional driver mutations, most commonly in *DNMT3A* (n=8), *TET2* (n=4) and *PPM1D* (n=3) (Fig.1c).

### Phylogeny of haematopoiesis and patterns of driver mutations

Phylogenetic trees were reconstructed from the presence or absence of SNVs across colonies (Extended Fig.1c, Methods). The phylogenetic trees of 3 patients with stable disease are shown in Fig.2 and for the remaining 7 patients in Fig.3 – these trees essentially depict the family relationships among cells currently contributing to blood production in each patient. Although the shapes and structures of the trees are unique to each patient, many common themes emerge. In patients with multiple driver mutations, the other drivers occurred both prior to or following *JAK2*^*V617F*^, as well as in independent HSCs, as previously reported^23–25^.

**Figure 2.**
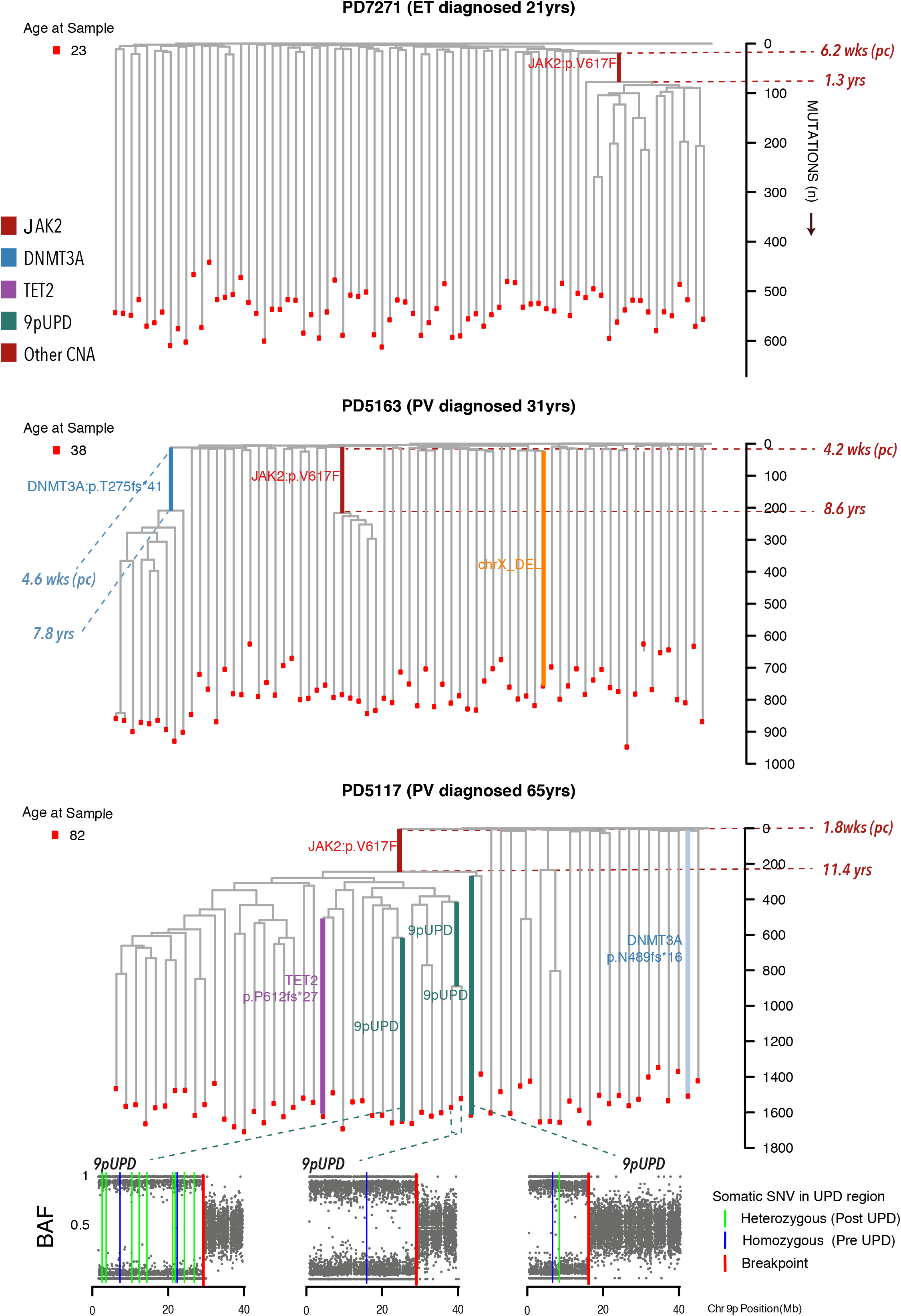
Phylogenetic histories of 3 patients with MPN driven by *JAK2*^*V617*^. The phylogenetic trees for 3 patients with stable *JAK2*^*V617F*^-mutated MPN diagnosed at different ages. PD7271, a 21 years old female, presented with asymptomatic isolated thrombocytosis in keeping with ET, and was treated with aspirin. PD5163, a 32 years old female, presented with splanchnic vein thrombosis, relatively normal blood count parameters, a raised red cell mass in keeping with PV, and was treated with Interferon-alpha. PD5117 was diagnosed with asymptomatic PV at age 64 on the basis of elevated blood counts and a red cell mass, and was treated with hydroxycarbamide. The tips of the branches represent individual colonies (red dots). Shared branches represent those mutations present across all downstream descendant colonies, and an end branch represents mutations unique to the single colony at its branch tip. Branch lengths are proportional to mutation counts shown on the vertical axes. Branches containing driver mutations and chromosomal aberrations are highlighted on the trees by colour. The corresponding times for the start and end of the shared branches harbouring driver mutations are shown on the trees. Ages at diagnosis and any progression of disease are shown in labelling above each tree, and ages at the time of sampling are shown to the left of trees. For branches with copy number aberrations, such as chromosome 9p uniparental disomy (UPD) in the phylogenetic tree of PD5117, we show the B-allele frequency (BAF) plots of part of chromosome 9p to highlight the chromosome breakpoints (vertical red line) for each acquisition. Heterozygous SNVs are mutations that occur after 9pUPD (vertical green line), whereas homozygous SNVs (that are not germline SNVs) would have been heterozygous SNVs prior to the 9pUPD but become homozygous as a consequence of the UPD. Given a clone-specific mutation rate, the proportion of heterozygous to homozygous SNVs on 9p can broadly indicate the timing of the UPD event. In this case, the leftmost 9pUPD occurred prior to other 9pUPD events due to the greater number of heterozygous mutations that have accumulated since acquisition. ET, Essential Thrombocythemia; PV, Polycythemia Vera.

**Figure 3.**
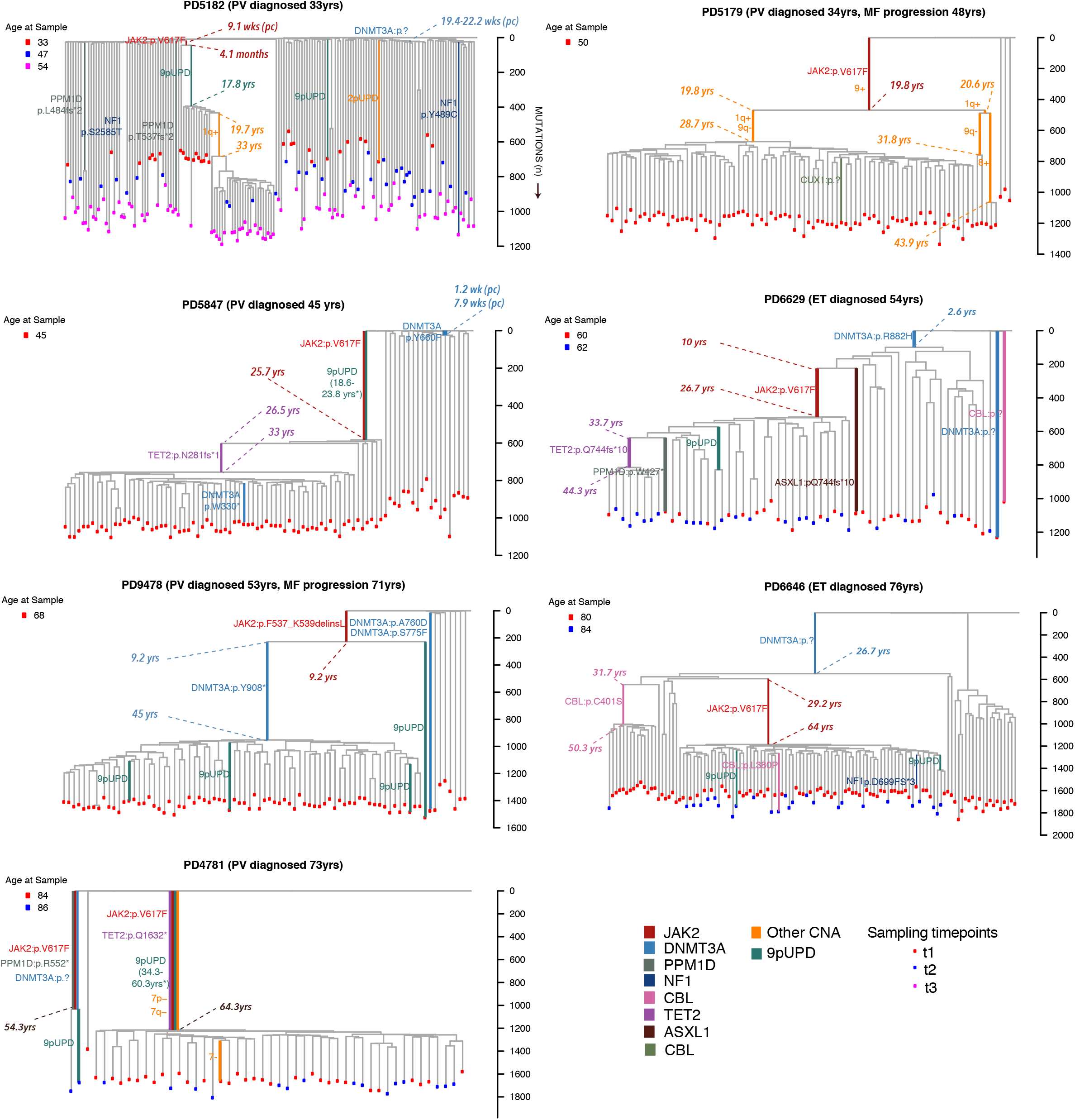
Phylogenetic histories of 7 patients with *JAK2*^*V617F*^-mutated MPN and clonal evolution. The phylogenetic trees of the remaining 7 patients with MPN who have evidence of multiple driver mutation led expansions. The vertical axis shows mutation counts. The tips of the branches represent individual colonies. Some patients were sampled at multiple timepoints, each timepoint highlighted by different coloured dots at branch ends. Age at diagnosis, times of any disease transformation, driver mutations and timing of mutations are depicted. *The timing of 9pUPD events in PD4781 and PD5847 are calculated using the proportion of heterozygous versus homozygous mutations on the UPD regions following estimations of clade-specific mutation rates. ET, essential thrombocythemia; PV, polycythemia vera; MF, myelofibrosis.

All patients had mixtures of some colonies with known driver mutations and some colonies without these drivers, suggesting that malignant haematopoiesis (both *JAK2-* and non-*JAK2* mutant clones) co-exist with normal blood production in MPN patients. The colonies without driver mutations shared few, if any, mutations with one another, evident as long, isolated branches in the phylogenetic trees – this demonstrates that the residual non-malignant blood production in MPN patients remains highly polyclonal, as seen in normal individuals^21^. Colonies with driver mutations typically shared tens to hundreds of mutations, including the driver mutation – this is evident as a ‘clade’ in the phylogenetic tree, namely a set of lineages descending from a shared ancestral branch, and confirms their clonal origin. Immediately beneath the shared branch containing the driver, we observe many short branches, each containing only tens of mutations – this represents a ‘clonal burst’ in which the original mutated HSC expands to a sizeable population of cells. The shortness of these initial branches implies that the clonal burst occurs rapidly – this is especially evident in the clones with multiple driver mutations (Fig.3).

We observed a number of instances in which similar genetic changes were acquired by unrelated clones, so-called ‘parallel evolution’. Chromosome (chr) 9p copy-neutral loss-of-heterozygosity due to uniparental disomy (UPD) was observed as multiple occurrences within the same patient (PD6646, PD5182, PD5117 and PD9478), often with unique breakpoints (PD5117 in Fig.2, Extended Fig.2a-e) as observed before^26^. Two separate acquisitions of chr1q+ and 9q-were noted in PD5179, affecting different parental chromosomes in each instance (Extended Fig.3a-c). Multiple mutations affecting the same oncogene were also observed within individual patients – 2-3 *DNMT3A* mutations in PD5847, PD6629, and PD9478; 2 *CBL* mutations in PD6646, and 2 *NF1* and *PPM1D* mutations in PD5182. (Fig.3). Many of these driver mutations were not part of the MPN clone (defined as the lineage harbouring mutated-*JAK2*). We also noted two independent acquisitions of *JAK2*^*V617F*^ in PD4781 (Fig.3) on different parental chromosomes (Extended Fig.3d-e). Taken together, the parallel evolution of similar genetic aberrations within patients suggests that there are patient-specific factors shaping the evolutionary trajectories of MPNs.

We did not identify novel coding or non-coding driver genes in our patients but noted a clonal expansion in PD6646, aged 80, with no known cancer-associated driver mutation in the ancestral shared branch (Fig.3 Extended Table 3). We also noted smaller wildtype expansions in PD5117, who was 82 years old (Fig.2). These findings provide evidence of the single cell origin of previous reports of CH lacking driver mutations 27, although what drives these clonal expansions and how they relate to old age remain unclear.

### Mutation acquisition in adulthood and impact of driver mutations

The number of somatic mutations in individual colonies was corrected for the size of the sequenced genome (affected by both copy-number aberrations and gender) and depth of sequencing. This adjusted SNV burden strongly correlated with patient age (Fig.4a) in keeping with a constant background rate of mutation acquisition throughout life^2,21,22^. Using a Bayesian approach, we modelled clade-specific background mutation rates in individual patients and accounted for excess SNV accumulation in early life due to rapid proliferation (Methods)^28^. In wildtype lineages, the median mutation acquisition rate was 17.9 per year (95% confidence interval (CI) 17.2-18.6, Fig.4b). Our estimates were confirmed by orthogonal mixed-effect modelling (Methods), and are consistent with previous studies^18,19^.

**Figure 4.**
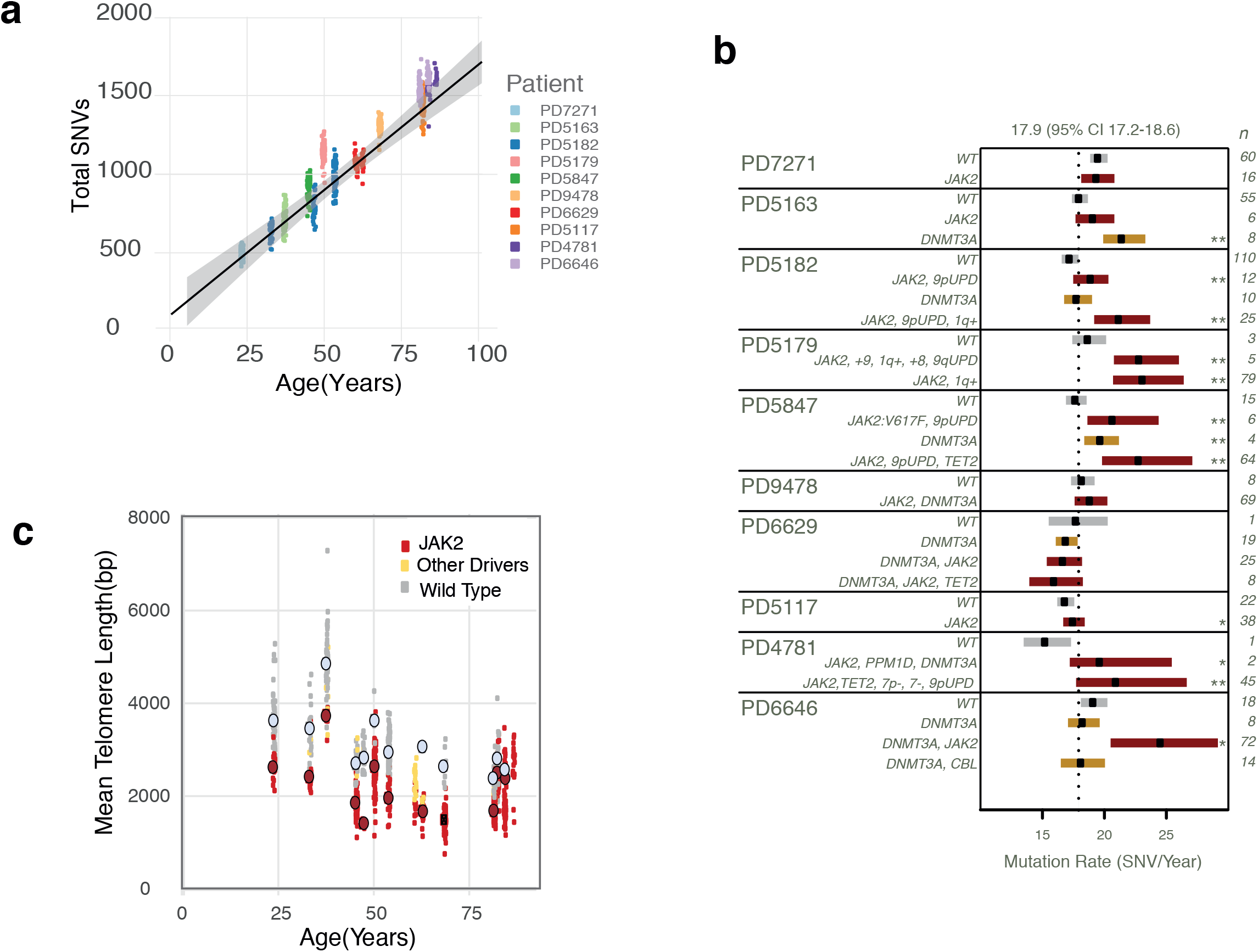
Mutation rates and impact of driver mutations. A. Total single nucleotide variants (SNV) and relationship to age. Dots represent single colonies that underwent whole genome sequencing and colours represent individual patients. Total SNVs represent non-germline SNVs adjusted for depth of sequencing. The black line shows the regression line and grey shading shows the 95% confidence interval. B. Clade specific mutation rates across individual patients. Patients and genotypes of clades are shown on the left. WT, wildtype clades are shown in grey bars, *JAK2*-mutated clades are shown in red and other mutant clades are shown in yellow. Number of colonies within each clade is shown on the right. The cohort wide estimate for the mutation rate in WT colonies is shown by the dotted black vertical line. C. Relationship between mean telomere length and age, for wildtype (grey dots), *JAK2*-mutated (red dots) and other mutant colonies (yellow dots). P-values *<0.5, **<0.01, ***<0.001 with multiple hypothesis correction.

In 7 of 10 patients, the mutant clades exhibited a significantly elevated mutation burden (1.5-5.5 more mutations/year) compared to matched wildtype counterparts (Fig.4b), most noticeably for *JAK2*-mutated clades. This increase could reflect greater background mutagenesis or increased cell division rates. Telomeres, the protective sequences at chromosome ends that progressively shorten with age and cell division, were significantly shorter in *JAK2*-mutated colonies (p=0.002, Fig.4c). Given that telomere lengths in individual mutant colonies are not fully independent measures due to their shared clonal origin, we corrected for phylogenetic distance, and still found that *JAK2*-mutated colonies had shorter telomeres (−864bps, CI 679-1035bp, p<0.001, Methods, Extended Fig.4). There was also a significant increase in C>T transitions at CpG dinucleotides in *JAK2*-mutated clades compared with wild-type colonies (Extended Table 4, Extended Fig.5) compatible with their increased cell division history^3^. We used the tops of the trees to capture the earliest stages of life and estimated the number of mutations that are acquired during cell division, as has been previously reported^21^. From our estimate that 1.7 mutations are acquired per cell division (Extended Fig.6), we calculated that mutant HSCs undergo an additional 0.7-3.9 cell divisions per year relative to wildtype counterparts to account for their higher mutation burden.

### Early acquisition of driver mutations during the lifetime of patients with MPN

Using these mutation rates, we then calculated estimates for the age at which driver mutations occurred in MPN patients. Specifically, we estimate the patient’s age at the beginning and end of branches containing driver mutations, as depicted on phylogenetic trees (Fig.2,3, Extended Fig.7) – this provides an age range within which the driver mutation or copy-number aberration most plausibly occurred. Copy-number aberrations were also independently timed by assessing the proportion of heterozygous and homozygous SNVs in affected regions, assuming clade-specific mutation rates (Methods).

In 5 patients in whom mutated-*JAK2* was the first driver event in the MPN clade, mutated-*JAK2* was acquired very early in life. PD5182, diagnosed at age 32yrs, acquired *JAK2*^*V617F*^ between 9.1 weeks post conception (pc) and 4.1 months age (CI 4.2 weeks (pc)-1.3yrs). PD7271, diagnosed at age 20.8yrs, acquired *JAK2*^*V617F*^ between 6.2 weeks (pc)-1.3yrs (CI 1 week (pc)-2.2yrs). The remaining 3 patients acquired mutated-*JAK2* during childhood at the latest by 8.6yrs (CI around latest age estimate of 7.3-10.1yrs, PD5163), 9.2yrs (7.7-10.8yrs, PD9478) and 11.4yrs (9.1-12.4yrs, PD5117). In these *JAK2-*‘first’ patients, the mean latency between mutated-*JAK2* acquisition and MPN diagnosis was 34yrs (range 20-54yrs). In a further two patients, *JAK2*^*V617F*^ occurred as the second driver within a mutated-*DNMT3A* clade, with disease latencies of 12.1yrs (PD6646) and 27.4yrs (PD6629) from *JAK2*^*V617F*^ acquisition. Latency to disease diagnosis from mutated-*JAK2* acquisition, irrespective of the ordering, was 31yrs (range 12-54yrs). In the remaining three patients (PD4781, PD5847, PD5179), we were unable to precisely time *JAK2*^*V617F*^ due to the presence of additional driver events, eg., 9pUPD, on the same branch. However, timing estimates of 9pUPD acquisition fell to decades before disease presentation (Fig.3) implying that *JAK2*^*V617F*^ acquisition occurred even earlier than this.

Mutations in *DNMT3A*, the gene most commonly detected later in life in the context of age related CH^13–15^, were also acquired *in utero* or childhood (Fig.3). PD5182 acquired *DNMT3A*^*ess.splice*^ between 19.4-22.2 weeks (pc) (CI 5.8 weeks (pc)-3.8 months), precisely the 20^th^, 21^st^ or 22^nd^ mutation from the start of life in that lineage. In PD5847, *DNMT3A*^*Y660F*^ was already acquired by the 23^rd^ mutation, which was between 1.2-7.9 weeks (pc) (CI 4 days (pc)-12.8 weeks gestation). The canonical mutation *DNMT3A*^*R882H*^ was acquired by 2.6yrs (CI 1.6-3.8yrs) in PD6629, and PD5163 acquired *DNMT3A*^*T275fs*41*^ by 7.8yrs (CI 6.5-9.5yrs). Acquisition of subsequent drivers was common in patients and separated by decades (Fig.3).

### Clonal expansion rates are variable and influence disease latency

The pattern of branching, specifically, the timing of ‘coalescences’ in the mutant clade, as well as the final clonal fraction reached, reflect the rate of clonal expansion from the time of driver mutation acquisition to the time of sampling. We modelled HSC population dynamics with a forward-time simulator using a continuous birth-death process. We used approximate Bayesian computation to generate patient- and clone-specific estimates of the selective advantage, or fitness, of mutant clones, that is, the additional proportion (*S*) by which each clone expanded per year (Fig.5a, Extended Table 5a and Fig.8, Methods).

**Figure 5.**
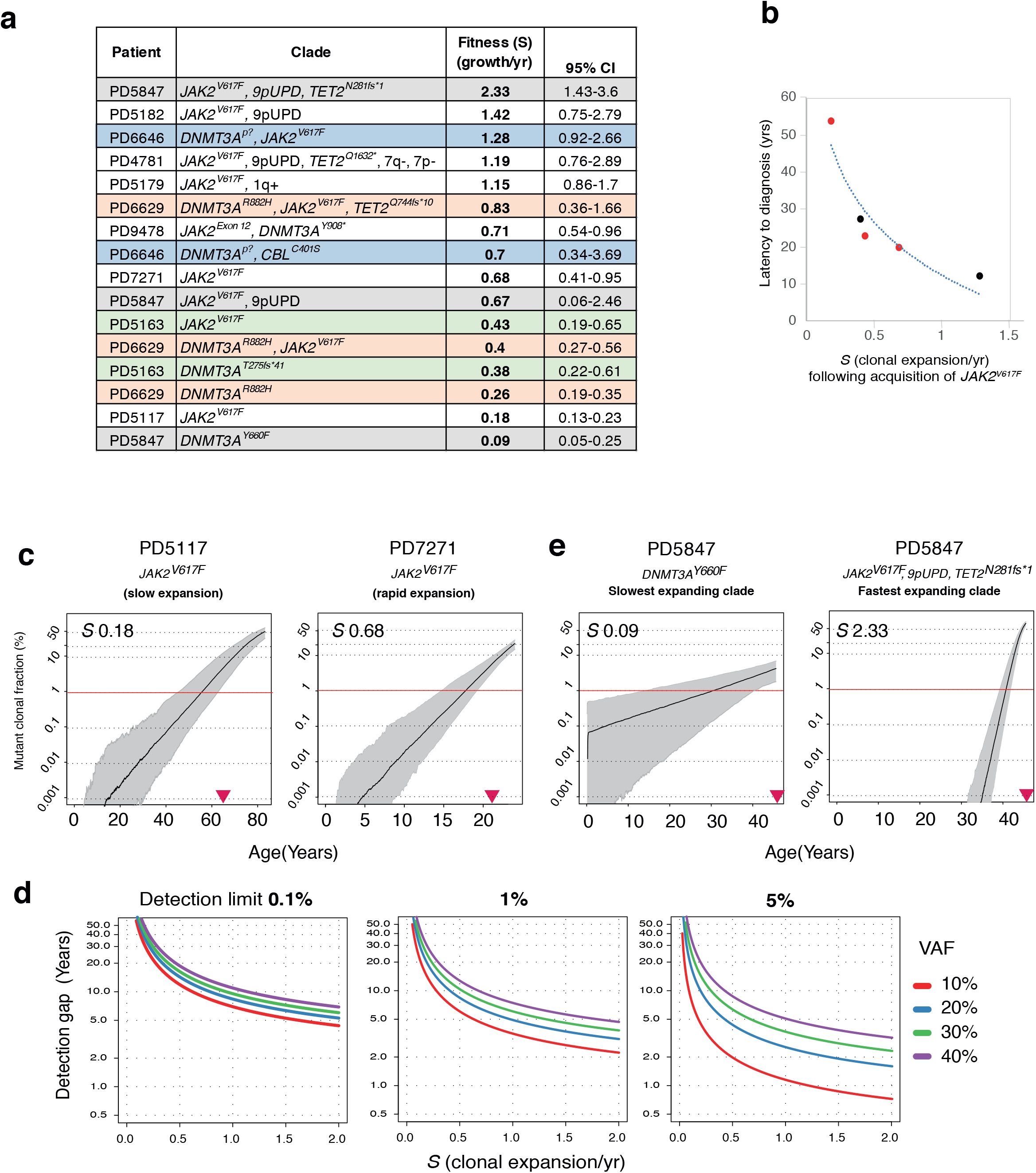
Clonal fitness and early detection. A. We define the fitness of clones by the selection coefficient, *S,* as the degree of clonal expansion occurring every year. *S* = 1 implies 100% additional growth, whereas S = 0 implies no change in clone size. The table shows *S* for clades across the cohort, ranked from highest to lowest, along with 95% confidence intervals (CI). *S* is highest for multiply mutated clades (2.33/year), and lowest for driver mutations common in clonal haematopoiesis (0.09/year). Coloured shading of rows are individual patients harbouring several different clades. B. The latency to diagnosis in relation to *S* following acquisition of mutated-*JAK2* is shown for 5 patients (PD7271, PD5163, PD5117, PD6646 and PD6629). Red dots represent patients with only mutated-*JAK2* as the driver mutation. Black dots represent *JAK2* mutation acquisition following mutated-*DNMT3A* C. The lowest and highest *S* in the context of the single driver mutation *JAK2*^*V617F*^, demonstrating the changing clonal fractions over the life of the patients. D. The modelled relationship between *S,* final variant allele fraction at MPN diagnosis, and the detection gap in years, assuming assay sensitivities of 0.1%, 1% and 5%. E. The lowest and highest *S* in the cohort detected in the same individual, from a very slowly growing *in utero* acquired mutated-*DNMT3A* clone, to a multiply mutated rapidly growing MPN clone. Pink arrowheads show age at diagnosis.

We observed variable rates of clonal expansion following acquisition of *JAK2*^*V617F*^ (Fig.5a). PD7271, the youngest MPN patient in our cohort, had a rapidly expanding *JAK2*^*V617F*^ clone at *S*=0.68/yr (CI 0.41-0.95). However, PD5117, one of the eldest patients in the cohort with diagnosis 54yrs after *JAK2*^*V617F*^ acquisition, had a much slower growing clone (*S*=0.18, CI 0.13-0.23, Fig.5a), not far above recent growth rates estimated for *JAK2*^V617F^-CH^29^. When *JAK2*^*V617F*^ was acquired on a mutated-*DNMT3A* backdrop, rates of expansion remained variable with *S=*1.28 (*JAK2*^*V617F*^/*DNMT3A*^*p.?*^ clone in PD6646, CI 0.92-2.66) and *S*=0.40 (*JAK2*^*V617F*^/*DNMT3A*^*R882H*^ clone in PD6629, CI 0.27-0.56, Fig.5a). Overall, rates of clonal expansion following the acquisition of *JAK2*^*V617F*^ were inversely proportional to the latency between mutation acquisition and disease diagnosis (Fig 5b).

Additional driver mutation acquisition corresponded with more rapid clonal growth. Within individual patients, we observed successive increases in expansion rates with sequential driver mutation acquisition (PD6629 *DNMT3A*^*R882H*^ *S*=0.26 (CI 0.19-0.35); *DNMT3A*^*R882H*^/*JAK2*^*V617F*^ *S*=0.40 (CI 0.27-0.56); *DNMT3A*^*R882H*^/*JAK2*^*V617F*^/*TET2*^*Q744fs*10*^ *S*=0.83 (CI 0.36-1.66), Fig.5a). The most rapidly growing clades in the cohort all harboured additional driver mutations, with *S* ranging from 1.28-2.33/yr (Fig.5a), which translates to the clone doubling in size every 7-10 months. We also observed very slow growing clones. In PD5847, the *DNMT3A*^*Y660F*^ clone was expanding at a rate of *S*=0.09 (CI 0.05-0.25, Fig.5a) since its acquisition in the first few weeks of life. This expansion rate is in line with selection estimates of *DNMT3A-*mutated CH from healthy aged individuals^29^ and demonstrates how mutation acquisition at the start of life, followed by a lifetime of slow expansion, can underlie CH observed in the elderly. Indeed, at very low selective coefficients, the probability of stochastic extinction of a clone was also higher, as would be expected (Extended Fig.9).

We considered whether selection bias for certain lineages during *in vitro* culture could have skewed our estimates of growth rates. Somatic mutations from the phylogenetic trees were deep sequenced using targeted recapture in bulk mature blood cells from the same patients (Fig. 1, Methods) to infer population estimates. We observed that clonal fractions of clades from phylogenetic trees were broadly concordant with population fractions measured in bulk blood samples (Extended Fig.10a). We estimated the proportion of lineages that may have diverged from shared branches harbouring driver mutations but which were not captured in our trees and also did not find any significant presence of lost lineages (Extended Fig.10b). This suggests that minimal sampling bias occurred from *in vitro* culture in our cohort, and also makes it unlikely that different clades are preferentially represented in different mature blood cell types, in accordance with previous observations^21^. We also used the population fractions of mutant clades in longitudinal bulk samples to corroborate the growth rates inferred from trees. We observed that clonal trajectories modelled from phylogenetic trees were in line with observed changes in clonal fractions in bulk longitudinal blood samples. Any differences in selected patients were explained by clinical interventions, in particular, treatment with interferon-alpha, either before or after sampling for phylogenetic trees (Extended Fig.11).

### Early detection of MPN driver mutations and rates of clonal expansion

Given the latency between mutation acquisition and clinical presentation, we asked how much earlier in life one might have detected the mutant clone prior to diagnosis. The growth trajectories for the different mutant clones were estimated from population simulations and correlated with patient age (Extended Table 5b). For the slowest growing *JAK2*^*V617F*^ clone (PD5117), we estimated that it took ~40 years for the mutant cell fraction to reach a 1% cell fraction, which was ~10 years before diagnosis. With detection limit of 0.01% cell fraction (1 in 10,000 cells), *JAK2*^*V617F*^ would have been detectable ~40 years before diagnosis (Fig.5c). With more rapid *JAK2*^*V617F*^ clonal expansion (PD7271), mutant cells would have been detectable at age 8yrs at 0.01% cell fraction (13 years prior to diagnosis, Fig.5c). Overall, with sensitive techniques detecting up to 1 in 10,000 aberrant cells^16,30^, we estimate it would have been possible to detect *JAK2*^*V617F*^ across all patients >10 to 40 years before disease presentation, irrespective of the timing of *JAK2*^*V617F*^ acquisition or the final clonal fraction reached (Fig.5d).

A complex and dynamic scenario can arise in patients with several mutant clades. In PD5847, clonal evolution has already occurred at diagnosis when the patient presented with life-threatening portal vein syndrome (Fig.3). In this patient, the *in utero* acquired *DNMT3A*^*Y660F*^ clone was growing modestly, taking ~30 years to reach a 1% clonal fraction. However, their *JAK2*^*V617F*^ clone then acquired both 9pUPD and *TET2*^*N281fs*1*^ which then rapidly expanded (*S=*2.33/year, Fig.5e) to reach clonal dominance within a decade prior to diagnosis. This illustrates the rapidity with which clonal progression can occur and emphasises the current unmet need for early detection of driver mutations and estimation of clonal trajectories for risk stratification of patients.

## Discussion

Our knowledge of the absolute timing of genomic aberrations and rates of expansion that drive human cancers is rudimentary. Sequencing of bulk cancer tissues, the final snapshots of a complex multi-step process of tumourigenesis, has captured the landscape of intra-tumour heterogeneity^31^, the relative ordering of genomic events^24,32,33^, and have provided broad estimates indicating that some chromosomal gains may occur several years to a decade or more before cancer presentation^4,5^. By using somatic mutations for lineage tracing, we re-trace life histories from early embryogenesis through to the single cell HSC origin of MPN and CH, and delineate driver mutation timing, clonal selection and clonal evolution over the life of 10 patients with MPN.

MPN are a common chronic blood cancer prevalent in up to 1 in every 1000-2000 individuals, with an increasing incidence with age^34,35^. Our study reveals, however, that regardless of age of diagnosis *JAK2*^*V617F*^ is usually acquired early in life (earliest appearance few weeks post conception). *DNMT3A* mutations, most commonly associated with age related CH^13–15^, were also acquired *in utero* and during childhood with slow and steady growth over life. Our 10 patients were not pre-selected other than to capture a broad range of ages, yet to find early driver mutation acquisition in all 10 consecutively sequenced patients makes it highly probable that this would be a feature of the majority of MPN patients. Our model of *in vivo* MPN development is also consistent with previous observations of *JAK2*^*V617F*^ detection prior to MPN diagnosis and in cord blood ^36,37^. Furthermore, the early acquisition of mutations associated with CH that slowly grow over life, provides an explanation for the very low burden CH detected in younger adults in some studies^16,30^. We identified that clonal expansions spanned the lifetime of MPN patients, often with sequential driver mutation acquisitions, including *JAK2*^*V617F*^ as a second driver event, as observed before^23^. The lifelong trajectories to MPN provide a new model for cancer development which may be relevant to other organs, given the abundance of mutations under selection across histologically normal tissues^7,9–11,13–16^, and the experimental tools developed here could be applied to other cancers.

MPNs are unique in that 40% of patients only harbour a single somatically acquired genomic event, eg. in *JAK2*^19^. However, we find the rate of clonal expansion upon *JAK2*^*V617F*^ acquisition to be variable, which may reflect differences in cytokine homeostasis, iron stores, bone marrow microenvironment, HSC lineage bias, inflammatory insults and germline influences^38–42^. Such factors may dynamically contribute over the lifetime of the individual. Factors influential in early life, due to altered embryonic HSC properties or stochastic drift, could also shape the future trajectories of nascent clones. Overall, growth rates associated with *JAK2*^*V617F*^ in MPN patients were greater than that inferred for *JAK2*^*V617F*^-CH^29^, which may account for the relative lack of clinical manifestations in the latter group. Rates of expansion for CH clones harbouring mutated-*DNMT3A* were 0.09-0.38/yr, in line with estimates from population based cohorts with CH^29^. We were unable to identify environmental or germline ‘triggers’ for either the start, or the speed of driver-mediated “clonal bursts” in our small cohort. Specifically, there were no prior exposures to chemotherapy. Concurrent potential exposures during life included smoking, obesity, infections (eg hepatitis C, mumps and whooping cough) and pregnancy.

MPN diagnosis is currently defined phenotypically, by blood count parameters and bone marrow histomorphology^43^. Our results indicate that the point at which a clinical diagnosis is made, represents one time-point on a continuous trajectory of lifelong clonal outgrowth, at which blood counts have reached certain thresholds, or clinical complications have already occurred. Current diagnostic criteria do not best capture when patients begin to have *disease* as life-threatening thromboses often trigger diagnosis. Furthermore, diagnostic criteria do not capture when individuals begin to be *risk* as those harbouring *JAK2*^*V617F*^ in the general population have increased risk of thrombosis, altered blood count parameters and gravely increased risk of future MPN^44^. Our data show that mutant clones will generally have been present for 10 to 40 years before diagnosis and would have been detectable for much of this time using sensitive assays. Clonal fractions of 1%, the common cut-off used for population screening studies, already reflect decades of clonal outgrowth, and the rate of expansion of *JAK2*^*V617F*^ strongly influences latency to disease presentation, more so than age at *JAK2*^*V617F*^ acquisition or clonal fraction at diagnosis. Taken together, the key to early detection and prevention may lie in both detecting low burden mutant clones early *and* in establishing their rate of growth, by repeated sampling, to capture those individuals on a future trajectory to clinical complications. The cornerstone of MPN management currently is aimed at normalising blood counts and reducing risk of thrombotic or haemorrhagic events – such treatments are mostly safe and well-tolerated, and could be offered to individuals with high-risk molecular profiles. Our data also provide a strong rationale for the ongoing evaluation of measures^45–48^ that target the *JAK2*^*V617F*^ clone in order to curb clonal expansion and subsequent clonal evolution.

## Acknowledgements

We thank Cambridge Blood and Stem cell biobank, funded by the Cambridge Cancer Centre and Wellcome Trust Cambridge Stem Cell Institute, CASM and DNA pipelines for their assistance. We thank Sam Behjati and Claire Harrison for comments. The study was supported by Cancer Research UK (JN), EHA Research Award (JN), MPN Research Foundation (JN) and the Wellcome Trust (PJC, ARG). PJC is a Wellcome Trust Senior Clinical Fellow. Work in the ARG Lab is supported by the Wellcome Trust, Bloodwise, Cancer Research UK, the Kay Kendall Leukaemia Fund, and the Leukaemia and Lymphoma Society of America. JN is a CRUK Clinician Scientist fellow. We thank the patients for their participation in the study.

## Author contributions

JN, ARG and PJC conceived and directed the study. NW performed genomic, phylogenetic and population dynamics analyses with JN. JL assisted with signature and telomere analyses. LM assisted with low-input sequencing and mutation signature analysis. ALG assisted with clinical information. JN and EJB prepared samples. JN wrote the manuscript with input from coauthors. Authors reviewed and approved the manuscript.

## METHODS

No statistical methods were used to predetermine sample size. The experiments were not randomised and the investigators were not blinded to allocation during experiments and outcome assessment.

### Patients and Samples

Patients were selected from *JAK2-*mutated MPN patients attending Cambridge University NHS Trust Hospital, UK, that had undergone previous whole exome sequencing. Apart from ensuring a wide representation of ages, patients were chosen at random. In doing so, we found that we captured different MPN subtypes, clinical presentations varying from asymptomatic blood count abnormalities to life threatening thrombosis, different MPN therapies, stable and progressed disease, and a wide mutation spectrum. This allowed us to capture a broad cross-section of MPN. Peripheral blood and bone marrow samples were obtained from patients with myeloproliferative neoplasms attending Cambridge Universities NHS Trust following written informed consent and ethics committee approval. Recapture samples were peripheral blood derived granulocytes apart from two samples which were whole blood and peripheral blood derived mononuclear cells. Constitutional samples were obtained from either buccal or T-cell DNA.

### *In-vitro* colony culture and next generation sequencing

Peripheral blood mononuclear cells were isolated from patient blood or bone marrow samples, cultured for 14 days in MethoCult 4034 (Stemcell) and single erythroid haematopoietic colonies (burst forming unit-erythroid, BFU-E) were plucked and lysed in 50ul of RLT lysis buffer (Qiagen). Library preparation for whole genome sequencing used enzymatic fragmentation and the NEBNext Ultra II low input kit (NEB) with 150bp paired end on either Illumina HiSeqX or Novaseq machines. Reads were aligned to the human reference genome (NCBI build37) using BWA-MEM.

### Somatic variant identification, genotype matrix of samples and variant loci level filtering

Single nucleotide variants (SNV) were identified using CaVEMan^49^ for each colony by comparison with both a matched germline sample, as well as an unmatched normal sample. Short Insertions and deletions were called using cgpPindel^50^. Copy-number aberrations (CNA) were identified using a matched normal ASCAT analysis. The union of colony SNVs and Indels was taken and reads counted across all samples belonging to the patient (colonies, recapture samples, buccal and T-cells) using VAFCorrect. The genotype at each locus within each sample is either 1 (present), 0 (absent) or NA (unknown). We inferred the genotype in a depth sensitive manner. Using a binomial model, we inferred genotype as either mutant or wildtype. The genotype was set to the most likely of the two possible states provided one of the states is at least 20 times more likely than the other. Otherwise the genotype is set to missing (NA). The VAF is usually 0.5 for autosomal sites, but for Chromosomes X, Y and CNA sites, we conservatively set it to 1/ploidy. For loss-of-heterozygosity (LOH) sites the genotype is overridden and set to missing if it is originally 0. A germline SNV Filter allowed for somatic mutation variant identification in the presence of modest levels of contamination in the germline sample and removal of germline variants. Further loci were removed if within 10bp of an indel, 10bp of each other, where more than a fifth of the colonies have depth<6, where more than a fifth of colonies had a missing genotype; where all samples harboured the variants; where the total mutant read count was significantly less than 0.9*Expected VAF*total depth across all colonies that have genotype=1 (Binomial Test); where the total mutant read count was significantly less than 0.9*0.5*total depth across all colonies that had at least 2 mutant reads (Binomial Test); and where the total mutant read count at sites with genotype=0 was significantly greater than 0.01*Depth.

### Colony Level Filters

Colonies were initially removed if the CaVEMan detection sensitivity was below 60%. In addition, colonies were tested for cross-contamination with other colonies from the same patient after tree construction. We required that the mean VAF of variants from a colony that mapped to private branches were not significantly less than 0.45 (Bonferroni adjusted one-sided binomial test) as this is evidence of some cross-contamination. For colonies that were not part of the same clade, cross-contamination was tested for by identifying colonies that exhibited a VAF significantly more than 5% VAF on non-ancestral branches (Bonferroni adjusted one-sided binomial test).

### Construction of phylogenetic tree topology

We constructed trees using Maximum Parsimony with MPBoot^51^. The input for the method is alignments based on the genotype matrix with missing values. This method also enables the rapid generation of bootstrap trees that correspond to the Maximum Parsimony trees that would be obtained by resampling the genotypes columns with replacement. Only SNVs were used to infer the topology but both SNVs and Indels were subsequently assigned to the branches. We developed an Expectation Maximisation method (R package “treemut”) to soft assign mutations to tree and to simultaneously estimate branch length. Having estimated the maximum likelihood probability that each mutation belongs to each branch we then hard assign mutations for ease of interpretation. Simulations indicated that this approach did not exhibit obvious biases in branch length estimation vs true branch length and that using the edge length implied by the hard assignment has a minimal effect on the deviation from the true edge length. Driver mutations and CNAs were assigned to the corresponding branches of the tree. Independent CAN events were confirmed by distinctness of breakpoints and haplotypes.

### Branch Length Adjustment for SNV Calling Sensitivity

The length of private branches of low depth colonies are likely to be underestimated because of the limited sensitivity of the variant calling. The per colony sensitivity was estimated in a non-parametric fashion from the identified germline SNVs by measuring the proportion of germline sites that were called by CaVEMan and private branches were scaled by 1/sensitivity. A similar approach is taken for shared branches where the sensitivity is estimated as the proportion of germline sites in which at least one of the colonies that share the branch has the variant called.

### Testing genes under selection

Genes that exhibit a deviation of ratio of non-synonymous to synonymous variants were evidence of non-neutrality. The SNVs and Indels were grouped by individual branches across the cohort and R package “dndscv” was used.

### Targeted sequencing of bulk peripheral blood (recapture) samples

We used Agilent SureDesign to design a baitset that captured all unmasked shared mutations. For private branches, we assayed up to 4 randomly selected unmasked variants per year of approximate branch length, capping the total number per private branch at 80. Variants that passed the SureDesign “most stringent” filter were preferentially selected. Sequencing on Illumina Novaseq was undertaken to depth of roughly 300-400x across all recapture samples.

### Mutational signatures

The SNVs that were mapped to the 10 patient trees were divided into per patient driver/wild-type clade-based groupings, where the mutations mapped to the clades were treated as an individual sample for the purposes of mutational signature extraction. *De Novo* Signature extraction was carried out using SigProfiler. Additional analysis was carried out using MutationalPatterns and custom scripts.

### Mutation rate estimations

We estimated mutation burden in the wildtype cells using two methods. We used linear mixed effects modelling to estimate the wildtype mutation rate using mutation burdens adjusted for depth of sequencing and excluded CNA regions, with age as a random effect and depth as a fixed effect. Only the timepoint with the most wild-type colonies is included in the model (this only affects PD5182). This fitted model has intercept 98.3 (95% CI 19.2-177.5) and the per patient wild type mutation rate in mutations per year is drawn from a distribution with mean 16.5 (95% CI: 14.8-18.2) and a variance of 1.1. The second method uses Bayesian modelling to jointly fit wild type rates, mutant rates and absolute time branch lengths under the assumption that the observed branch lengths are Poisson distributed with *Mean* = *Duration* × *Sensitivity* × *Mutation Rate*. The model incorporates an excess mutation rate in early life that will add an average of 33.5 mutations during the first 6 months post conception. The mean branch timings are directly sampled from the posterior distribution and by construction the resulting trees are guaranteed to have a root to tip distance that matches the sampling age of the colony. Models were fitted across four chains each with 20,000 iterations including 10,000 burn-in iterations. The model was coded in R and Rstan and inferred using the Rstan implementation of Stan’s No-U-Turn sampler variant of Hamiltonian Monte Carlo method^52^. The resulting posterior mean and standard error of each patient’s wild type rate were combined in a random effects meta-analysis using the R package metafor.

### Methods for timing LOH and amplifications

The timing methods require a rough approximation for the number of expected detectable mutations, *L*, in the LOH/CNA region for the duration of the branch. Firstly, we estimate a local relative somatic mutation rate for mutations detectable by CaVEMan in autosomal regions. The rate is measured by counting distinct mutations across a panel of samples consisting of those colonies in the 10 patients that do not exhibit copy number aberrations (350,371 mutations across 594 colonies). The genome is divided into 100Kb bins and the number of passed somatic mutations is counted across all samples in the panel, to give a count *c*_*i*_ for bin *b*_*i*_. The probability that a given mutation occurs in bin *i* is estimated by 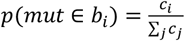 and with standard error in that estimate of 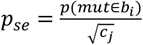 For a given copy number region H then 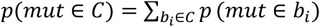. For a branch of duration *t* and with global mutation rate *λ* then *E*(*L*) = *λtp*(*mut*) ∈ *C*) and 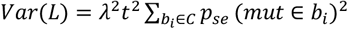 where it should be noted that errors in *t* and *λ* are not included in the variance estimation. All somatic mutations that occur prior to the LOH event, occurring a fraction *x* along the branch, will be homozygous with detection sensitivity *S*_*HOM*_ and those after will be heterozygous with detection sensitivity *S*_*HET*_. We model the mutations as arriving at a constant rate along the branch and fit the following model for *x* : *N*_*HET*_ ~ *Poission*(1 − *x*)*Ls*_*HET*_) and *N*_*HOM*_ ~ *Poission*(*xLs*_*HOM*_) with priors *x* ~ *Uniform*(0,1) and *L* ~ *N*(*E*(*L*, *Var*(*L*) and where *S*_*HOM*_ = 0.5 (assuming perfect detection of homozygous mutant variants) and *S*_*HET*_ is estimated from germline SNPs as previously discussed. Somatic mutations that occur prior to the CNA event, occurring a fraction N along the branch, have an equal chance of exhibiting VAF=1/3 or VAF=2/3, whereas those occurring after the event will always have VAF=1/3. 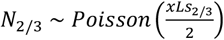 and 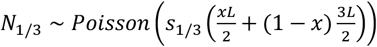 where the priors are as in the LOH model above. The detection sensitivities *s*_1/3_ and *s*_2/3_ are similar to *s*_*het*_ because of the additional sequencing depth afforded by the duplication. Unlike the LOH case, the value of *x* will be relatively unaffected by *L* because of the similarity of *s*_1/3_ and *s*_2/3_.

### Telomere analysis

The mean telomere lengths of the colonies were estimated using Telomerecat with batch correction using cohort wide information to correct the error in F2a counts as detailed. Given the slight discrepancies in length found on multiple readings, each telbam was analysed 10 times and the average length was used. Given that colonies sharing a driver mutation share a more recent common ancestor, and are therefore, not truly independent measures of telomere lengths, we measured telomeres adjusting for phylogenetic distance between colonies. To specifically ask how much shorter telomere lengths become as a result of a JAK2-mutation, we fit a phylogeny aware mixed model for the mean telomere length with a patient specific intercept using the MCMCglmm library in R (Iterations = 100001:1099001; Thinning interval = 1000; Sample size= 1000; DIC: 11187.69). This identifies an average loss of 21bp of telomere length per year of life and that *JAK2*-mutant clades exhibit ~864bp shorter telomeres.

### Continuous Time Birth Death Model

Each cell has a rate of symmetric division and a rate of symmetric differentiation (or death). Asymmetric divisions do not affect the HSC genealogy and are therefore not explicitly included in the model. Let l be the wild type rate of symmetric division, measured in divisions per day. We model selective advantage *s* as an increased rate of symmetric division *α*_*mut*_ = *α*(1 + *s*). We assume during the growth phase that the cells population grows unrestrained by death. Once the specified equilibrium population size, *N*, is reached then the death rate *β*, which is the same for every cell, matches the average division rate. Following the acquisition of a driver the mutant cell population grows stochastically until the population is sufficiently large, when the growth becomes essentially deterministic following a logistic growth function where in the early stages the exponential growth process exhibits an annual rate of growth *S*, given by: *S* = *exp*(*αs*) − 1. The above model is implemented using the Gillespie algorithm where the waiting time until the next event is exponentially distributed with a rate given by the total division rate + total death rate, this event is then division with probability=total division rate/(total division rate + total death rate). If the event is Division then the choice of which cell is given by a probability proportional to the cell’s division rate whereas if the event is Death then all cells are equally likely to be chosen. Implementation was in C++ with an R based wrapper. The simulator maintains a genealogy of the extant cells, together with a record of the number of symmetric divisions on each branch, the absolute timing of any acquired drivers and the absolute timings of branch start and end. The package also provides mechanisms for sub-setting simulated genealogies whilst preserving the above per branch information.

### Approximate Bayesian Computation

Approximate Bayesian Computation (ABC) was used to reproduce the timing and shape of clonal expansions in our phylogenetic trees using our simulator to generate sampled trees. The procedure was as follows: Infer mutation rate, *λ*, as the mean root to tip distance divided by the age of sampling. The observed mutation count at the start and end of the branch carrying the driver in the experimental tree is denoted by *M*_*start*_ and *M*_*end*_ respectively. Fix symmetric division rate at 1 division per year. Sample N from *Log*10(*N*) ~ *Uniform*(3,6.5). Sample age of driver acquisition by resampling mutation counts from a Poisson distribution: 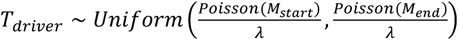. Sample S from *S* ~ *Uniform*(0.05,5). Simulate the population: Simulate the tree with initial division rate of 0.1 per day until population has grown to the equilibrium population size. Simulate neutral evolution until time *T*_*driver*_. Save the state of the simulation (*). Introduce the driver with the specified selection coefficient. If the driver lineage dies out before the sampling age is reached then return to the saved state (*) and try again. A tree with the observed number of mutant samples is subsampled from the population of extant cells. For each simulation the following summary statistics were calculated (i) Total deviation in the simulation mutant clade’s number of lineages through time (LTT) with respect to the patient clade of interest, and (ii) deviation of the simulation-based population clonal fraction with respect to the patient clade’s aberrant cell fraction. In all cases but two the aberrant cell fraction was calculated as the proportion of sampled mutant clades. For PD5163_JAK2 and PD5182_JAK2 the aberrant cell fraction at diagnosis is used, prior to the commencement of interferon-alpha treatment which led to clone size reduction. The total distance score is then calculated as a combination of the above two scores. For each clade between 726,678 and 959,453 simulations were run. In each case the posterior distribution was approximated by the top 0.02% simulations.

### Aberrant Cell Fraction

In recapture samples, a per branch aberrant cell fraction can be calculated as twice the aggregate mutant read fraction where only autosomal variants that map to the branch and are outside of the copy number aberrant regions are included. For each patient the trajectory of the aberrant cell fraction was retrieved from the top 0.2% of simulations. The simulator periodically takes snapshots at approximately daily intervals and measures the aberrant cell fraction as the current fraction of cells that carry the driver mutation. Subsequently, for each simulation these daily snapshots are binned into a common sequence of 10-day periods over each of which the aberrant cell fraction is averaged. We present the 2.5%, 50% and 97.5% quantiles for each binned period calculated across the simulations.

### Stochastic Extinction

The probability of extinction in a homogenous birth death process is the ratio of death rate to birth rate^53^ which in our case is 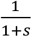 or equivalently 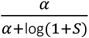. In the ABC simulations we record the number of attempts required to introduce the drivers. We then verified that the simulator behaved as expected by gathering all simulations across the analysed mutant clades and then restricted to the 13,048,861 simulations where the driver was introduced after development (>1 year post conception). The simulations were binned into Selection Coefficient bins of width=0.05 and log10(N) bins of width=0.1. The extinction probability was estimated in each bin using the maximum likelihood estimator 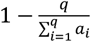 where à is the number of simulations in the bin and each simulation, *i*, fixes after *a*_*i*_ attempts.

### Start of Life Polytomies

The polytomies were used to estimate lower and upper bounds for the mutation rate per symmetric division during embryogenesis^21^, whereby the number of edges with zero mutation count at the top of the tree (up to the first 10 mutations) is inferred from the number and degree of polytomies assuming an underlying tree with binary bifurcations. The mutations per division are assumed to be Poisson distributed. A maximum likelihood range is then calculated in two steps first using the 95% confidence interval of the proportion, *p*, of zero length edges and with this leading to a maximum likelihood estimate for the Poisson rate as −log *p*.

---

**Extended Figure 1.**
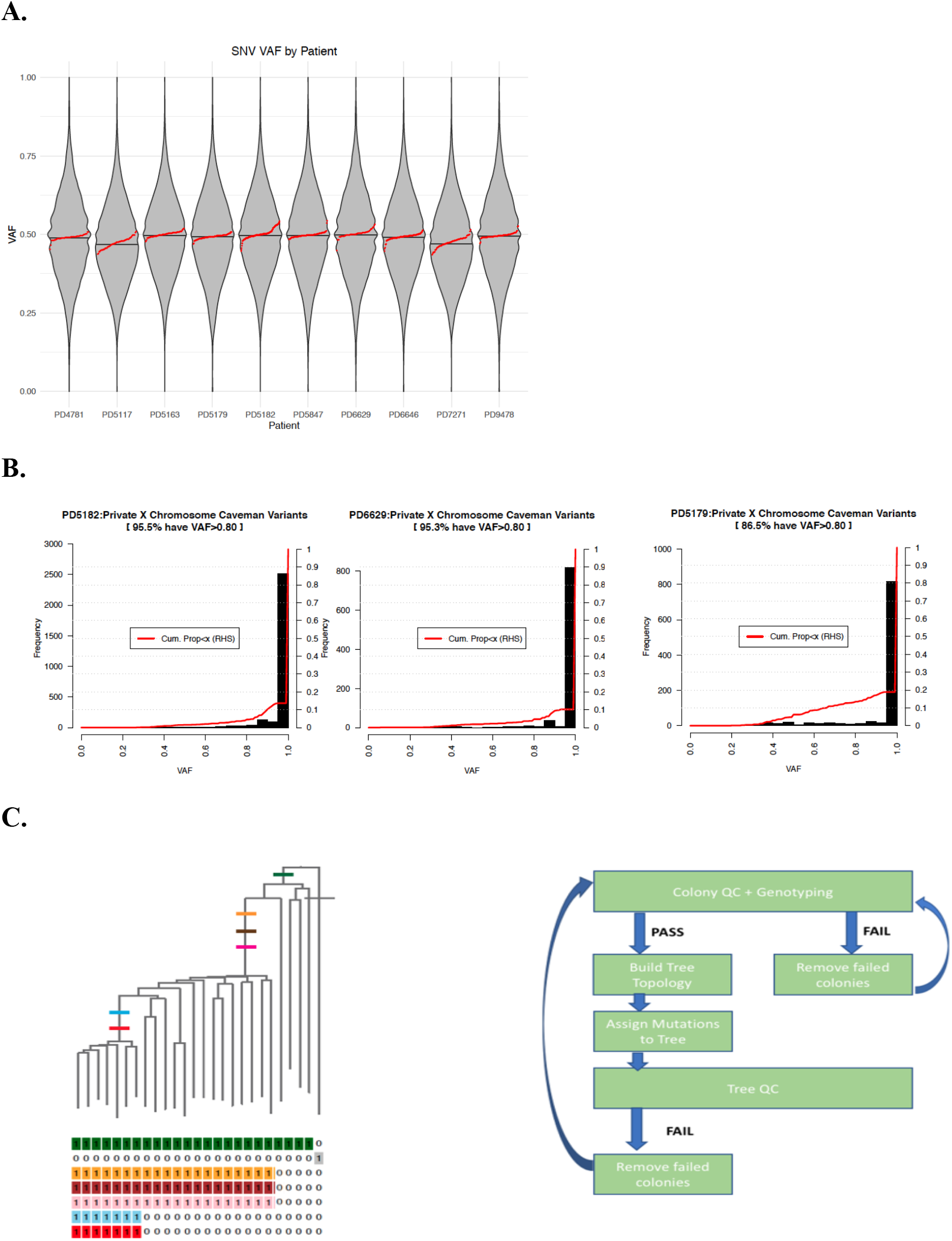
Using somatic mutations from clonal samples to build phylogenetic trees. **A.** The per patient VAF distribution pooled across colonies. The mean VAF of individual colonies is shown as red dots. Only autosomal SNVs are shown, excluding those in regions with copy-number aberrations and loss-of-heterozygosity. The plot shows that the colony VAFs are close to 0.5. **B.** VAF of variants on Chr X in male patients. The black bars show that only a minority of mutations are subclonal and potentially associated with acquisition during *in vitro* culture. The red line shows the cumulative proportion of chromosome X variants with a VAF less than the x axis threshold. **C.** Model of a phylogenetic tree on the left constructed using the presence or absence of mutations across the colonies, as shown below the tree. On the right, we depict the broad process of phylogenetic tree building once somatic mutations have been called. QC, quality control. Genotyping refers to the assignment of mutations to individual colonies, as present, absent or unknown.

**Extended Figure 2.**
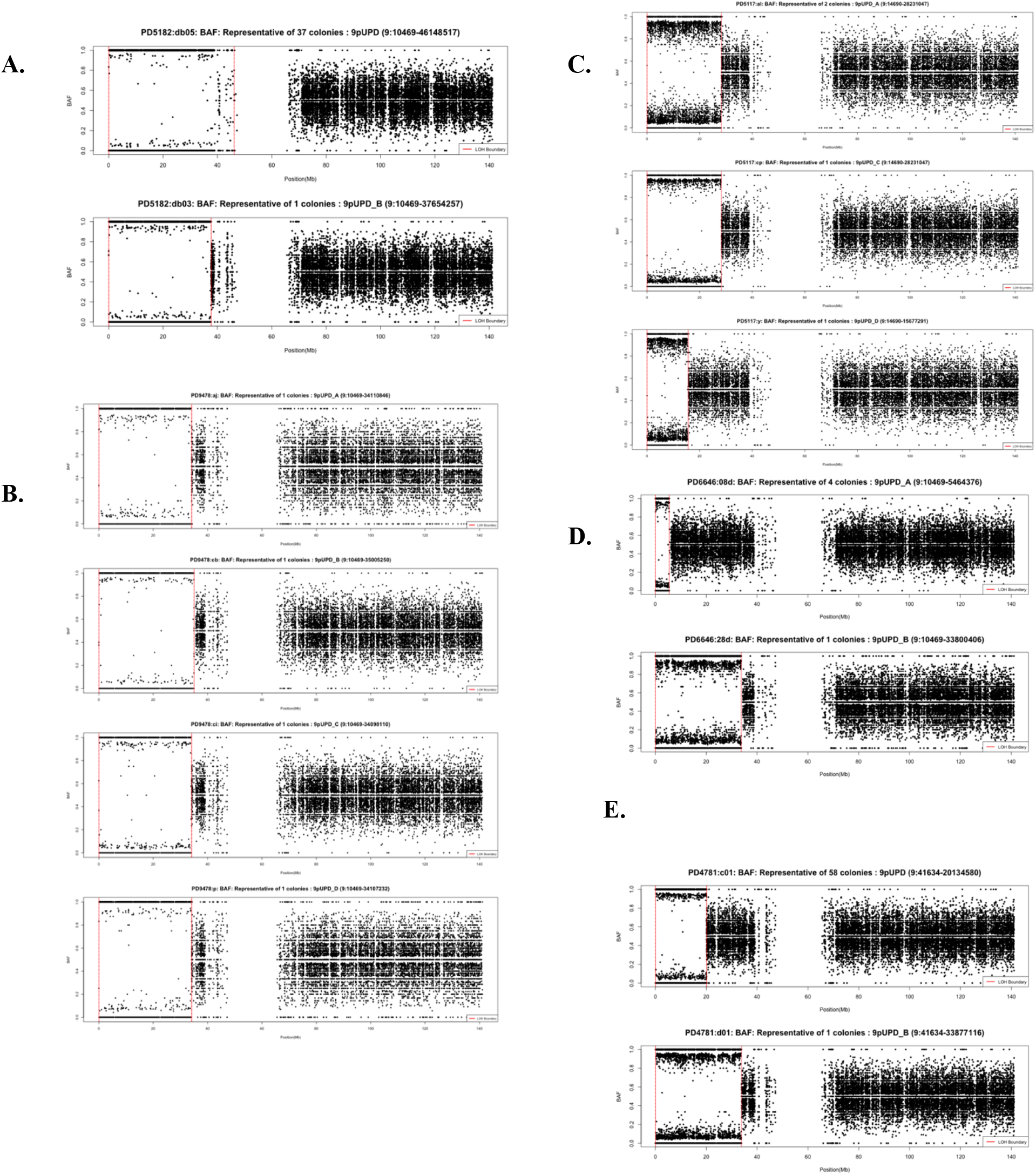
Parallel evolution observed in phylogenetic trees. B-allele frequency plots showing the regions of 9pUPD in different clades within the same patient. The vertical red lines show the boundaries of the LOH. 9pUPD events have distinct breakpoints in PD5182 (A). PD9478 9pUPD events have similar but distinct breakpoints (B). In PD5117 the top two events have the same breakpoints upon close examination of germline polymorphisms in the region, whereas the lower event is distinctly different (C). The PD6646 9pUPD events have very distinct breakpoints (D). The two 9pUPD events in PD4781 have distinct breakpoints (E). These events involve UPD of different paternal chromosomes each having acquired *JAK2*^*V617F*^ acquisition independently.

**Extended Figure 3.**
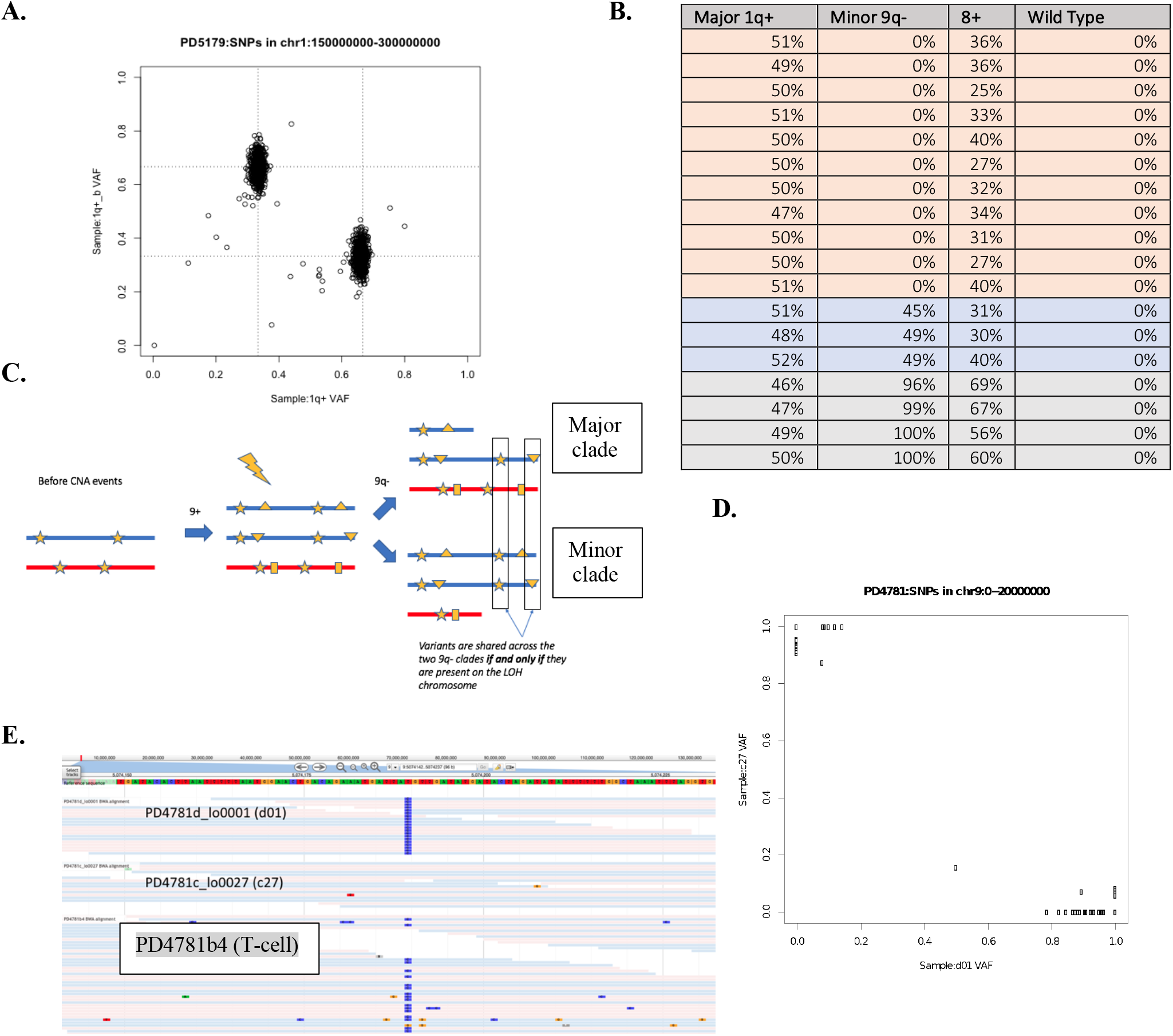
Multiple acquisitions of 1q+ and 9q-in PD5179 and *JAK2*^*V617F*^ in PD4781. **A** The aggregate VAF of SNPs in the 1q+ region for samples in the 1q+ major clade versus 1q+ minor clade in PD5179. SNPs at a VAF of two-thirds in one clade are at one-third in the minor clade, and vice-versa, confirming that different parental chromosomes are amplified in each clade. **B.** The chr 9q variants that map to the JAK2/9+ ancestral branch exhibit a clear pattern in the VAF. Samples in the major clone have VAF=0.5, in keeping with the loss of one copy of the amplified parental chromosome. Where samples in 9q-have VAF=0, the samples in the 8+ clade have VAF=1/3 and where the samples in 9q-have VAF=1, the samples in the 8+clade have VAF=2/3. This confirms the sequence of events in **C**, showing 2 independent acquisitions of 9q-with different parental chromosomes being lost in the major and minor 1q+ clades. **D.** SNP VAF analysis in PD4781 samples shows that 9pUPD involves a different parental chromosome in each instance. SNPs that have a VAF ~1 for 9pUPD samples in the JAK2-mutant dominant clade have a VAF ~0 in 9pUPD samples in the JAK2-mutant minor clade. **E.** PD4781 is heterozygous for the 46/1 haplotype as seen by the lower panel DNA reads from rs12343867 locus. Above this, sample c27, from the dominant *JAK2*-mutant clade, is now wildtype for the 46/1 haplotype as a result of 9pUPD, but sample d01, from the *JAK2*-mutant minor clade, is now homozygous for 46/1.

**Extended Figure 4.**
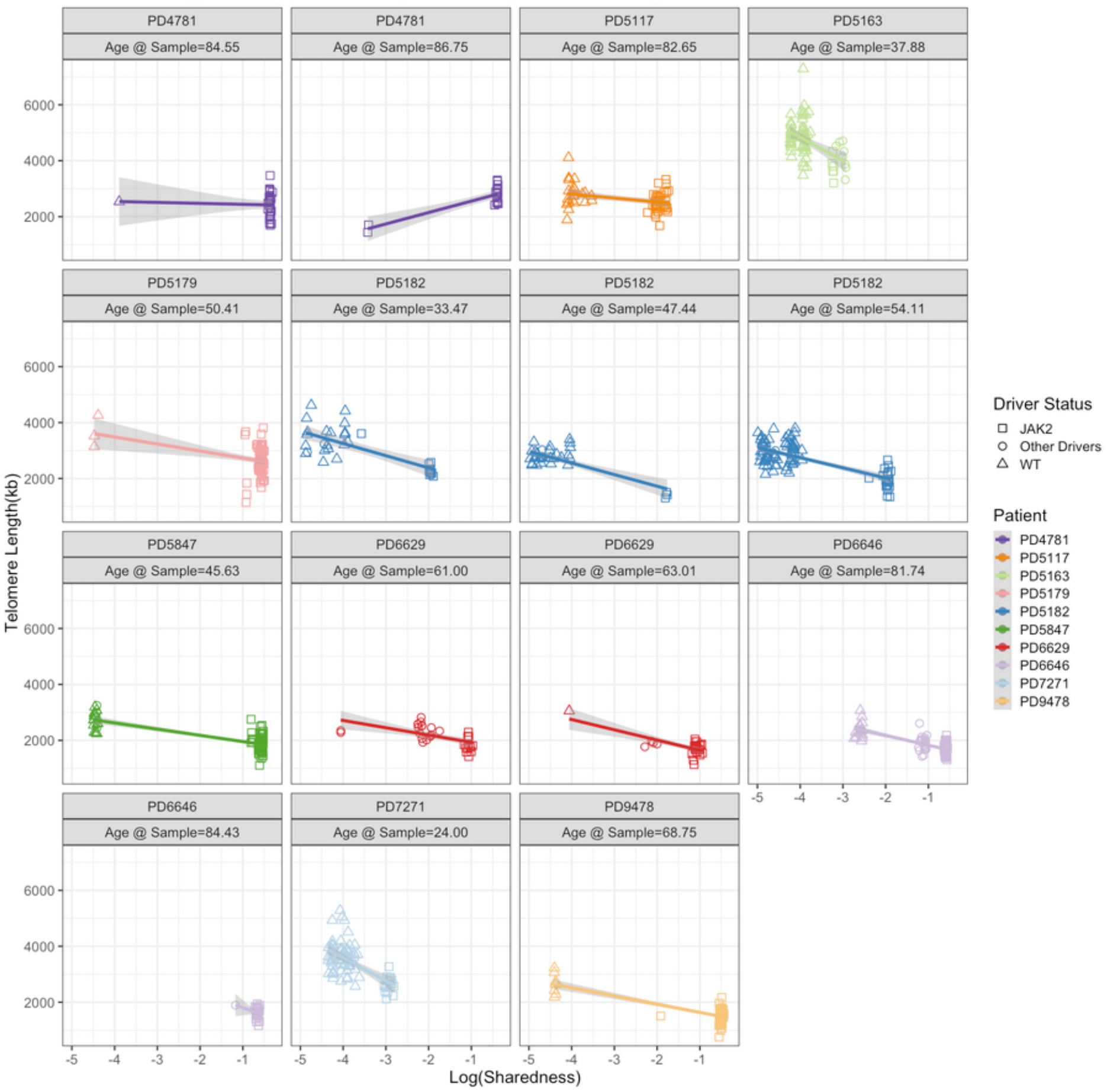
Phylogenetically aware telomere analysis. Following the observation that *JAK2* mutant colonies had significantly shorter telomere lengths than wild type colonies, or colonies with other driver mutations, we controlled for the fact that *JAK2*-mutant colonies have a more recent shared ancestor, and therefore, the measures within an individual patient are not independent of one another. We defined ‘sharedness’, that captures the degree of shared lineage history as a weighted average of the proportion of sampled clones that share each mutation. The figure above shows that telomeres do shorten in line with increased phylogenetic ‘sharedness’ in keeping with the increased cell divisions during clonal expansion.

**Extended Figure 5.**
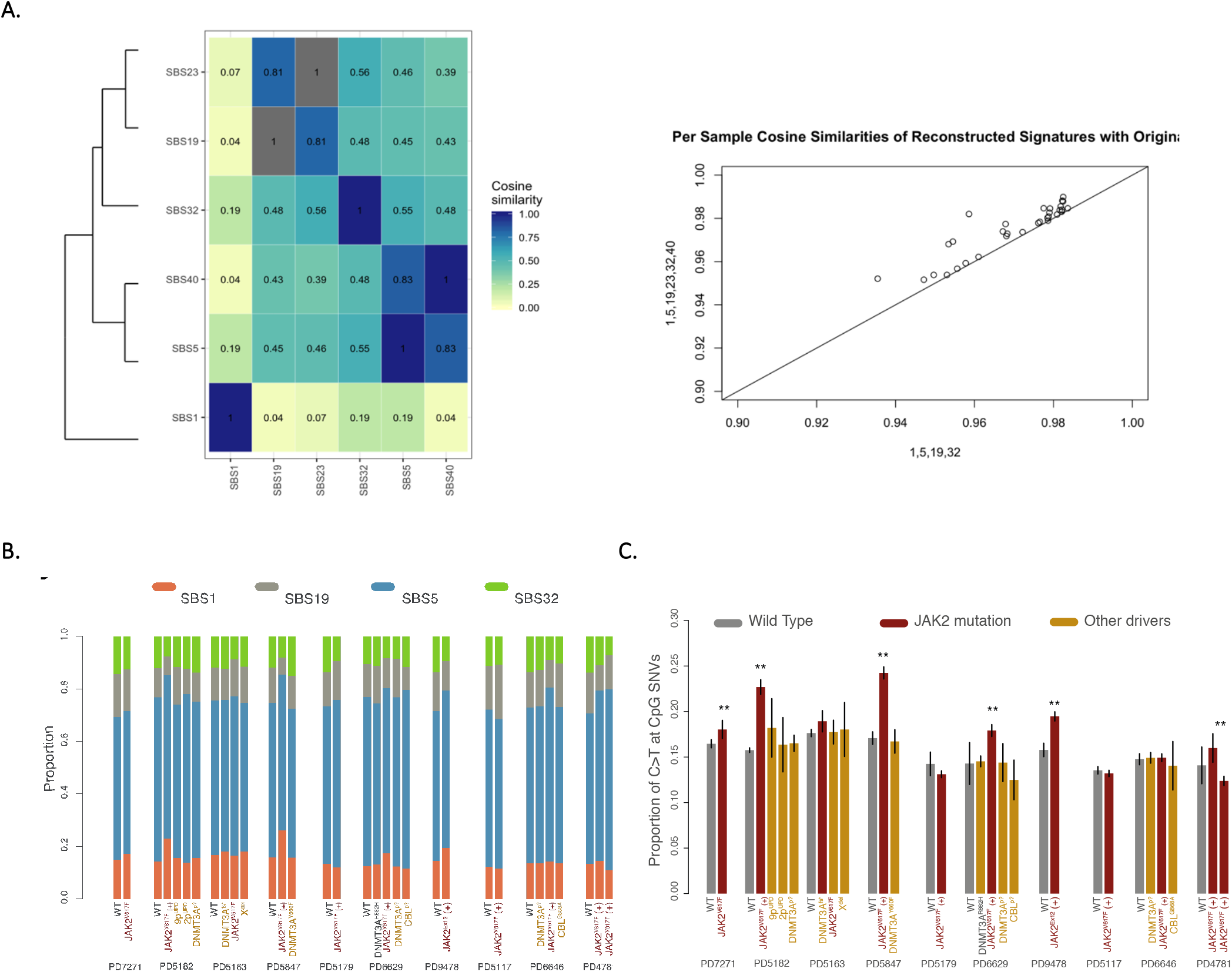
Mutational signatures. No novel signatures were discovered in addition to the standard PCAWG 60 signatures^54^. The identified signatures were SBS1, SBS5, SBS19, SBS23, SBS32 and SBS40. **A.** Comparison reduced signature set (SBS1, SBS5, SB19 and SBS32) versus the set (SBS1, SBS5, SBS19, SBS23, SBS32 and SBS40). Pair SB19 and SB23 had a high cosine similarity (0.81) as did SBS5 and SBS40 (0.83) as shown in the left panel. Removal of SBS23 and SBS40 resulted in an acceptable loss in reconstruction accuracy (mean cosine similarity 0.970 vs 0.975) as shown on the right. **B.** Signature contributions of SBS1, SBS5, SBS19 and SBS32 on a per-patient/per-clade basis. Single base substitution mutational signature 5 (SBS5), thought to represent a time-dependent mutational process active in all tissues, was the predominant mutational process in colonies accounting for 61% of SNVs (258,573 mutations). **C.** The proportion of C>T transitions at CpG dinucleotides across WT, *JAK2*-mutated and colonies with other driver mutations. ** p<0.01

**Extended Figure 6.**
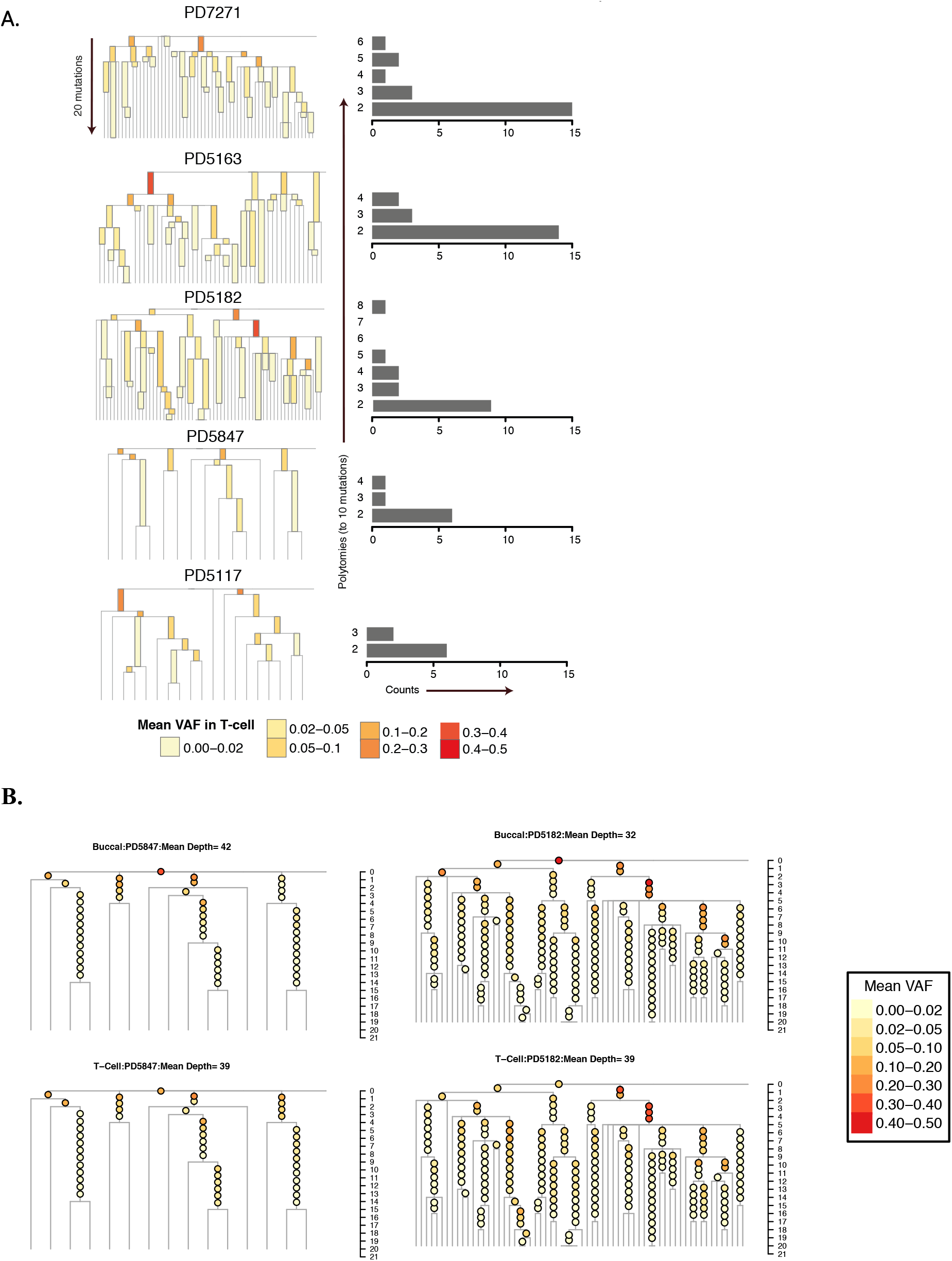
Somatic mutations during embryonic development. We used the pattern of branch splits at the tops of the phylogenetic trees to infer the mutation rate per cell division during early life, as described previously^21^. **(A)** above shows the top segments of phylogenetic trees, up to 20 mutations of molecular time, from five patients with adequate (>10 wildtype lineages) diversity at the top of their trees. Yellow to red shading shows the corresponding variant allele fraction (VAF) in bulk T-cells that underwent whole genome sequencing to an average depth of 38x. For PD7271, T-cells also underwent targeted recapture to higher depth of coverage of 385x. To the right of the expanded trees, the distribution of branch splits (2-way splits versus multiway polytomies) is shown and used to infer the mutation rate per symmetrical hematopoietic stem cell division. We observed a total of 228 lineages by 10 mutations of molecular time. Of the 227 symmetrical self-renewing cell divisions this would have required, 42 were mutationally silent, leading to a median estimate of 1.7 (range 1.4-2.1) mutations per cell division during early life. The rapid drop off in VAFs for somatic mutations from the early phylogenetic tree in bulk T-cell DNA is consistent with the early divergence of this tissue from the myeloid lineage. **B.** The top segments (up to 20 mutations of molecular time) of phylogenetic trees from two patients (PD5182, left; PD5847, right). Yellow to red shading shows the corresponding variant allele fraction (VAF) in buccal DNA (upper trees) and T-cells (lower trees). The mean depth of sequencing for buccal and T-cell samples is shown in the tree labels. We see very similar VAF distributions for early mutations from the phylogenetic trees in both buccal and T-cell DNA in two patients, suggesting that both these tissues diverge from the myeloid lineage early in life.

**Extended Figure 7.**
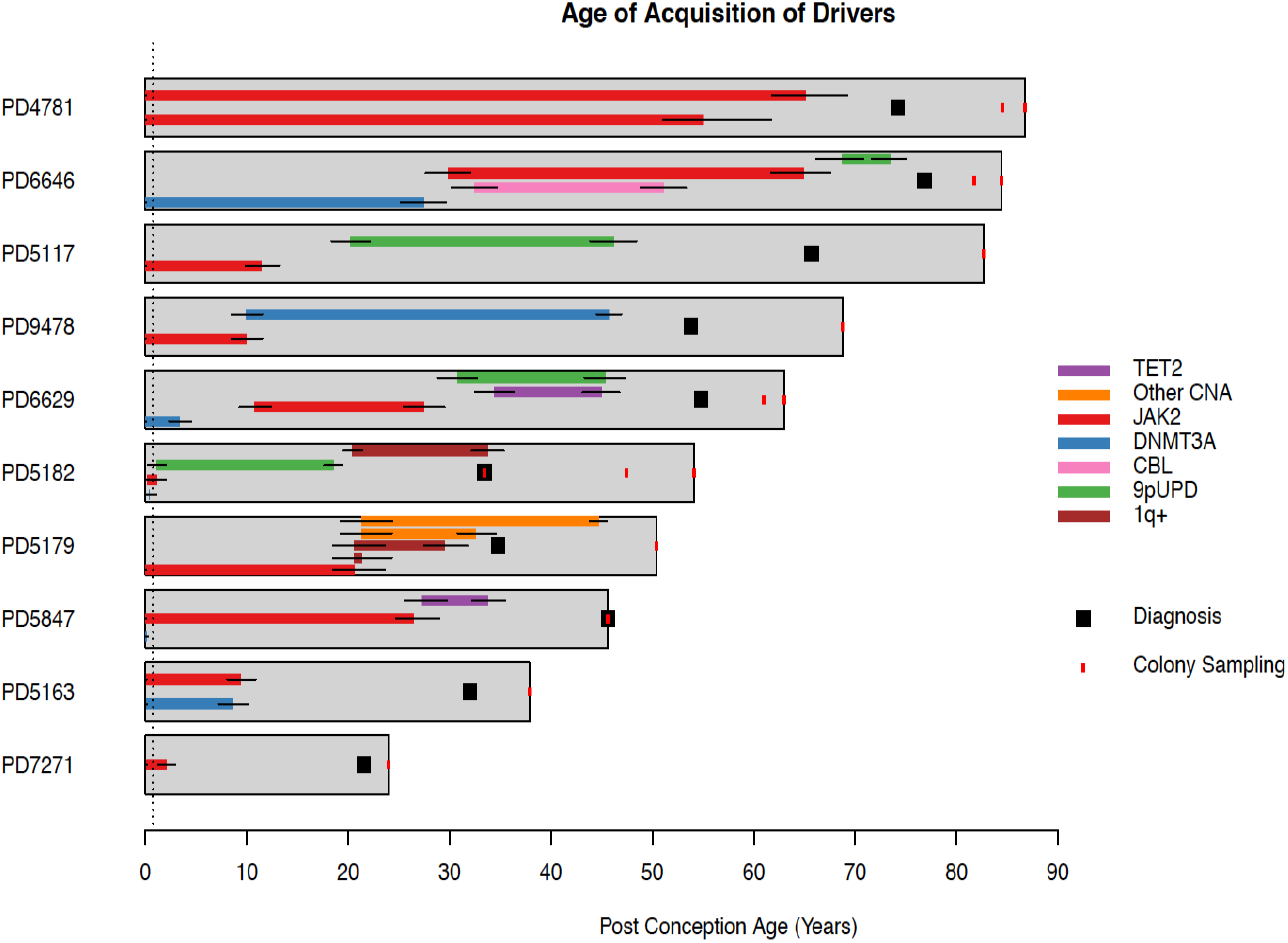
Summary of timing of driver mutation and CNA acquisition in patients. Each horizontal grey box represents an individual patient from the start of life until the last colony sampling timepoint. The x axis shows post conception age (which is age in years + 0.75). Within each grey box is shown the timing of driver mutation acquisition and copy number aberrations. The start and ends of each coloured box represent the median lower and upper bounds of time estimates corresponding to the start and end of the shared branches harbouring driver mutations. Black lines show the 95% credibility intervals for the start and end of the branches carrying the drivers.

**Extended Figure 8.**
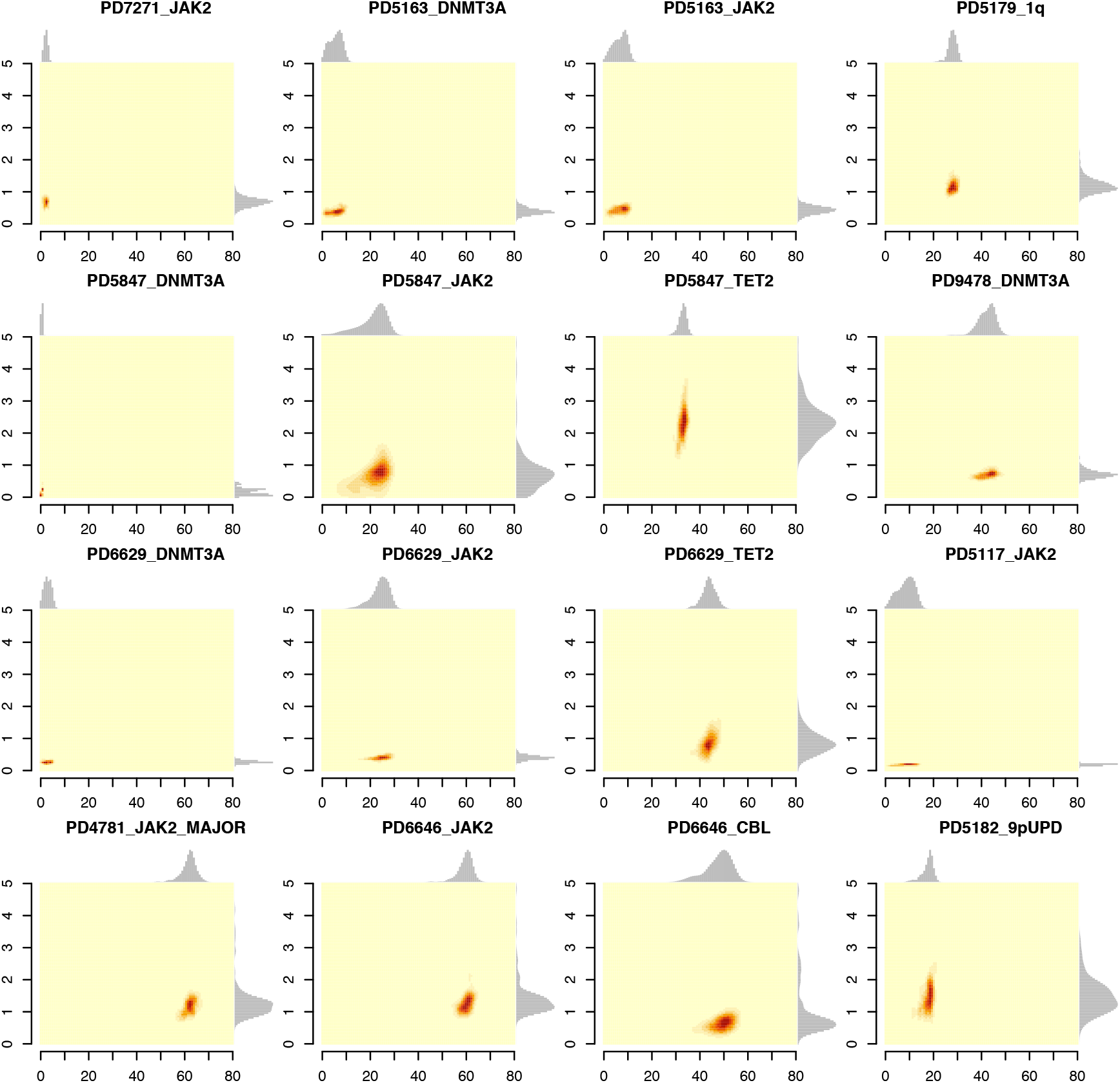
Posterior distribution of growth rates (S) from Approximate Bayesian Computation. The figure shows the smoothed posterior density distribution of the selection coefficient vs driver timing for all analysed clades. Marginal distributions are also shown. It is worth noting that the mass of the marginal selection coefficient distribution generally lies away from the edges of the prior distribution (0.05-5). The prior distribution for driver timing is clade dependent and is largely determined by the mutation count at the start and end of the associated branch. Incorporation of both clonal fractions and lineages through time as summary statistics in the approximate Bayesian computation allowed for narrower estimates of selection and could account for clades that were acquired very early in life with rapid growth, but did not reach a large clonal fraction later in life.

**Extended Figure 9.**
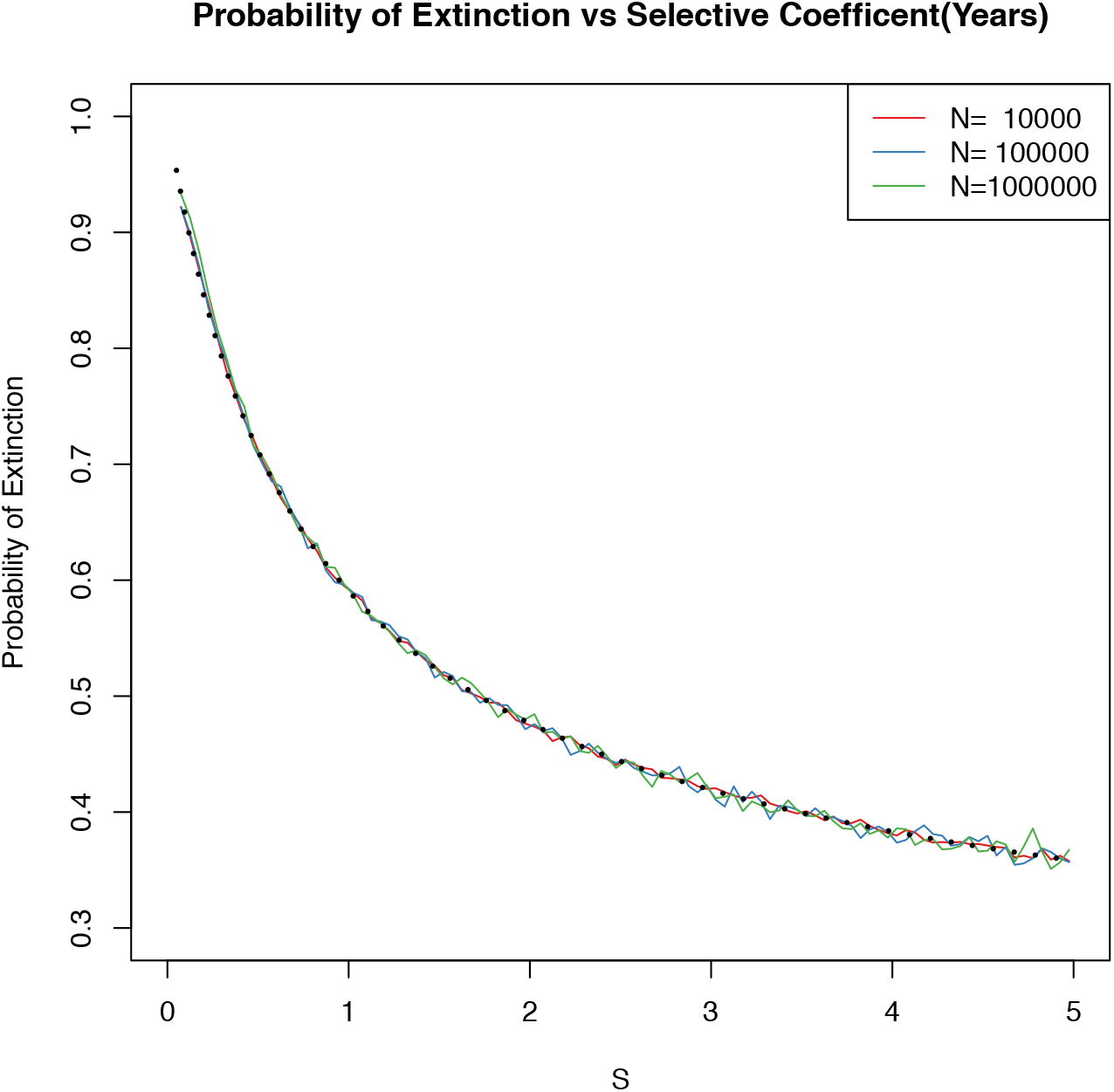
Probability of stochastic extinction for clones with different selective coefficients. The graph depicts the relationship between the selective coefficient (or fitness) of a driver mutation harbouring HSC and the likelihood of stochastic extinction after acquisition of the driver mutation. We recorded the number of attempts of driver mutation introduction across a total of ~13 million HSC simulations undertaken during the approximate Bayesian computation analysis. Simulations with driver acquisition>1 year post conception were binned into selection coefficient and total HSC population size bins. The empirical distribution of the number of driver mutation introduction attempts in each bin was converted into a bin specific maximum likelihood probability of extinction. The theoretical extinction probability (dotted line) is overlaid on the chart. S, selective coefficient (that is, the proportional increase in clone size per year) modelled as an increase in the rate of symmetrical HSC cell division due to a driver mutation.

**Extended Figure 10.**
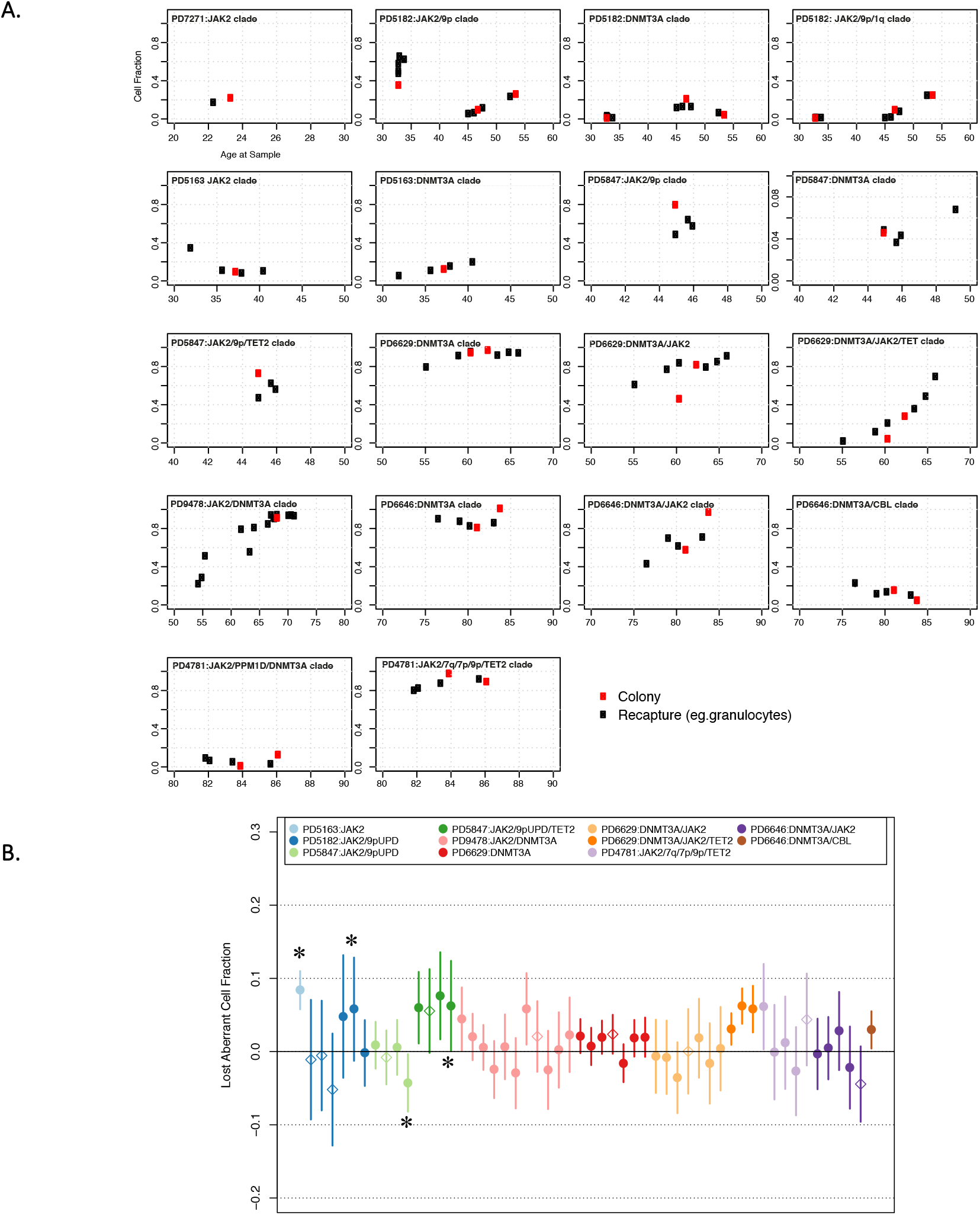
Mutant clonal fractions in colonies and bulk samples over time. **A.** Bulk samples (eg granulocytes) underwent targeted sequencing for mutations from the phylogenetic trees. In colonies, the aberrant cell fraction (ACF) is the clonal fraction as a proportion of all colonies (red dot). In bulk samples, the cell fraction is calculated as twice the mean VAF of variants that map to the shared ancestral branch of that clone on the phylogenetic tree (excluding variants that are in a copy-number aberrated/loss-of-heterozygosity regions) (black dots). The x-axis is patient age reflecting the different timepoints sampled. **B.** 95% confidence intervals for the difference in parent ACF and the aggregate of descendant daughter ACFs from phylogenetic tree clades. The confidence intervals are calculated assuming the sampling distribution of the aggregate mutant read fraction for each branch is approximately normally distributed. Diamonds indicate those recapture samples closest to the colony sampling and show no evidence of the presence of lineages in the population which were not captured in phylogenetic trees. *Samples taken whilst the patient was on Interferon-alpha therapy.

**Extended Figure 11.**
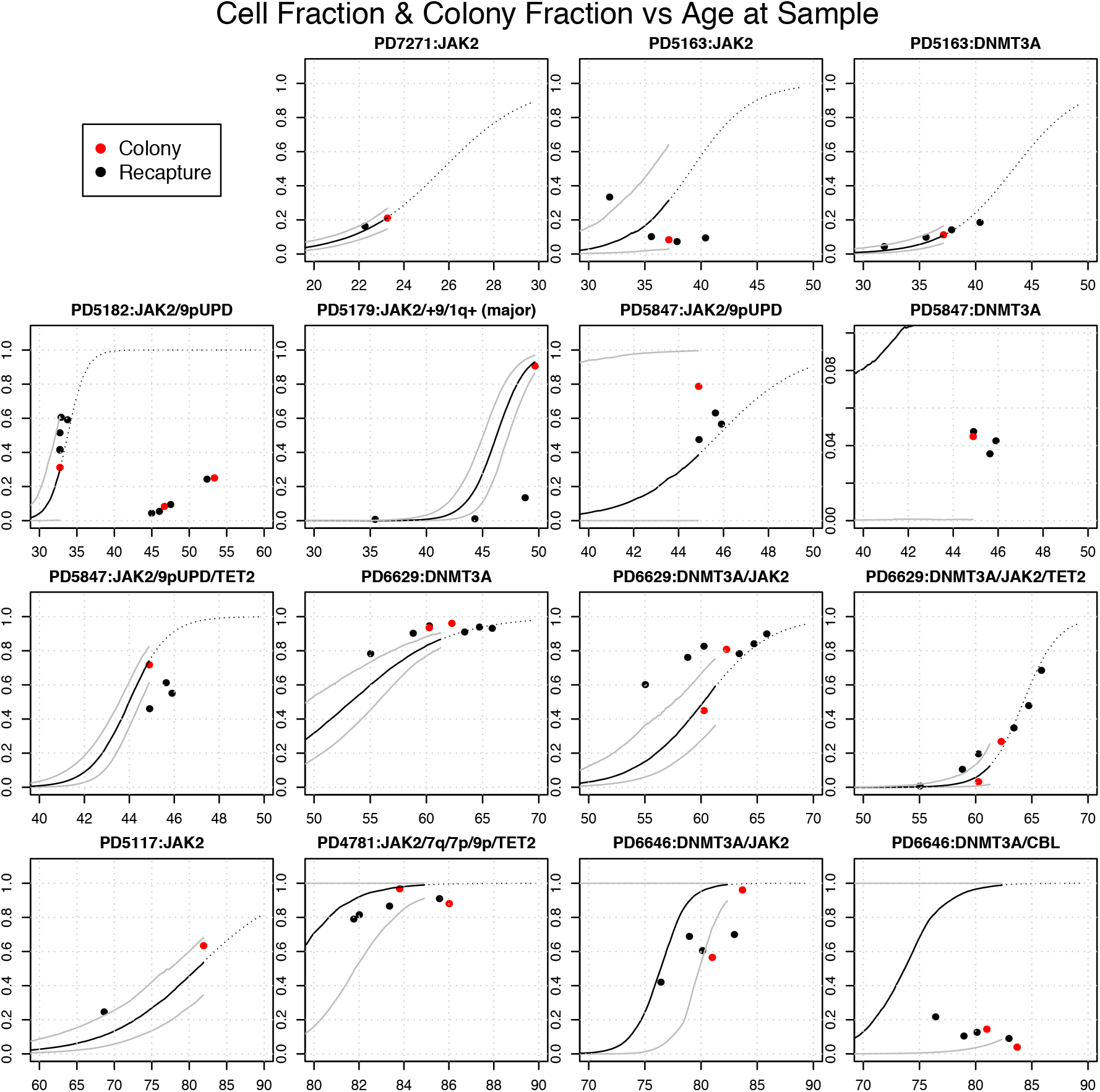
Comparison of aberrant cell fraction trajectories based on estimated *S* and aberrant cell fraction measurements from bulk samples. Bulk samples (eg granulocytes) underwent targeted sequencing for mutations from the phylogenetic trees. In colonies, the aberrant cell fraction (ACF) is the clonal fraction as a proportion of all colonies (red dot). In bulk samples, the cell fraction is calculated as twice the mean VAF of variants that map to the shared ancestral branch of that clone on the phylogenetic tree (excluding variants that are in a copy-number aberrated/loss-of-heterozygosity regions) (black dots). The x-axis is patient age reflecting the different timepoints sampling. Here, we overlay the aberrant cell fraction trajectories from the top 0.2% of simulations. Black lines show the median cell fractions and grey lines show 95% confidence bounds. The dotted line infers the future trajectory of growth of the clone beyond the sampling time of phylogenetic trees using the growth rate *S* and accounting for a sigmoid clonal trajectory as clonal dominance is approached.

**Extended Table 1.** Patient cohort and clinical characteristics.

| Patient | Diagnosis | Gender | Age at diagnosis, yrs | Blood counts: diagnosis (top), latest colony timepoint (bottom) |  |  |  | Cyto reduction | Thrombosis | WHO defined disease progression (age at progression) | Haematological response at time of colony sampling | Death (age at death or last follow up) |
| --- | --- | --- | --- | --- | --- | --- | --- | --- | --- | --- | --- | --- |
| | | | | Hb, g/L | Hct | WBC, $\times 10^9/L$ | Plts, $\times 10^9/L$ | | | | | |
| PD7271 | ET | F | 20 | 145 | 0.42 | 8.9 | 923 | None | N | N | NA | N (27) |
|  |  |  |  | 143 |  | 7.9 | 640 |  |  |  |  |  |
| PD5163 | PV | F | 32 | 164 | 0.49 | 11.6 | 435 | IFN (age 32-44) stopped due to cytopenia. | Y (portal / splenic vein) | N | Y | N (46) |
|  |  |  |  | 127 |  | 4.4 | 133 |  |  |  |  |  |
| PD5117 | PV | F | 65 | 166 | 0.50 | 10.6 | 831 | HC | N | N | Y | Y (89) |
|  |  |  |  | 136 |  | 4.1 | 271 |  |  |  |  |  |
| PD5182 | PV | M | 33 | 179 | 0.54 | 10.0 | 604 | IFN; became refractory aged 50, switched to HC | N | N | N | N (54) |
|  |  |  |  | 140 | 0.45 | 5.0 | 397 |  |  |  |  |  |
| PD5847 | PV | F | 44 | 190 | 0.59 | 20.0 | 504 | IFN | Y (portal vein) | N | N | N (51) |
|  |  |  |  | 190 | 0.59 | 20.0 | 504 |  |  |  |  |  |
| PD9478 | PV | F | 53 | 201 | 0.60 | 5.1 | 308 | None | N | PPV-MF (71) | N | N (76) |
|  |  |  |  | 140 |  | 15.9 | 111 |  |  |  |  |  |
| PD4781 | PV | F | 73 | 151 | 0.46* | 9.5 | 634 | HC | Y (TIA) | N (86) | N | Y (86) |
|  |  |  |  | 108 |  | 18.0 | 193 |  |  |  |  |  |
| PD6646 | ET | F | 77 | 145 | 0.44 | 7.7 | 804 | HC; Pipobroman | N | N (87) | N | Y (87) |
|  |  |  |  | 106 | 0.32 | 2.8 | 409 |  |  |  |  |  |
| PD6629 | ET | M | 54 | 139 | 0.43 | 13.5 | 842 | IFN, HC | Y (TIA) | N (70) | N | N |
|  |  |  |  | 149 |  | 8.7 | 433 |  |  |  |  |  |
| PD5179 | PV | M | 34 | 182 | 0.55 | 64.3 | 200 | HC; switched to ruxolitinib after MF transformation | Y (Budd-Chiari, CVA) | PPV-MF (48) | N | Y (51) |
|  |  |  |  | 98 |  | 12.5 | 646 |  |  |  |  |  |

\* PV diagnosed on red cell mass study
HC = hydroxycarbamide; IFN = interferon-alfa

**Extended Table 2.**
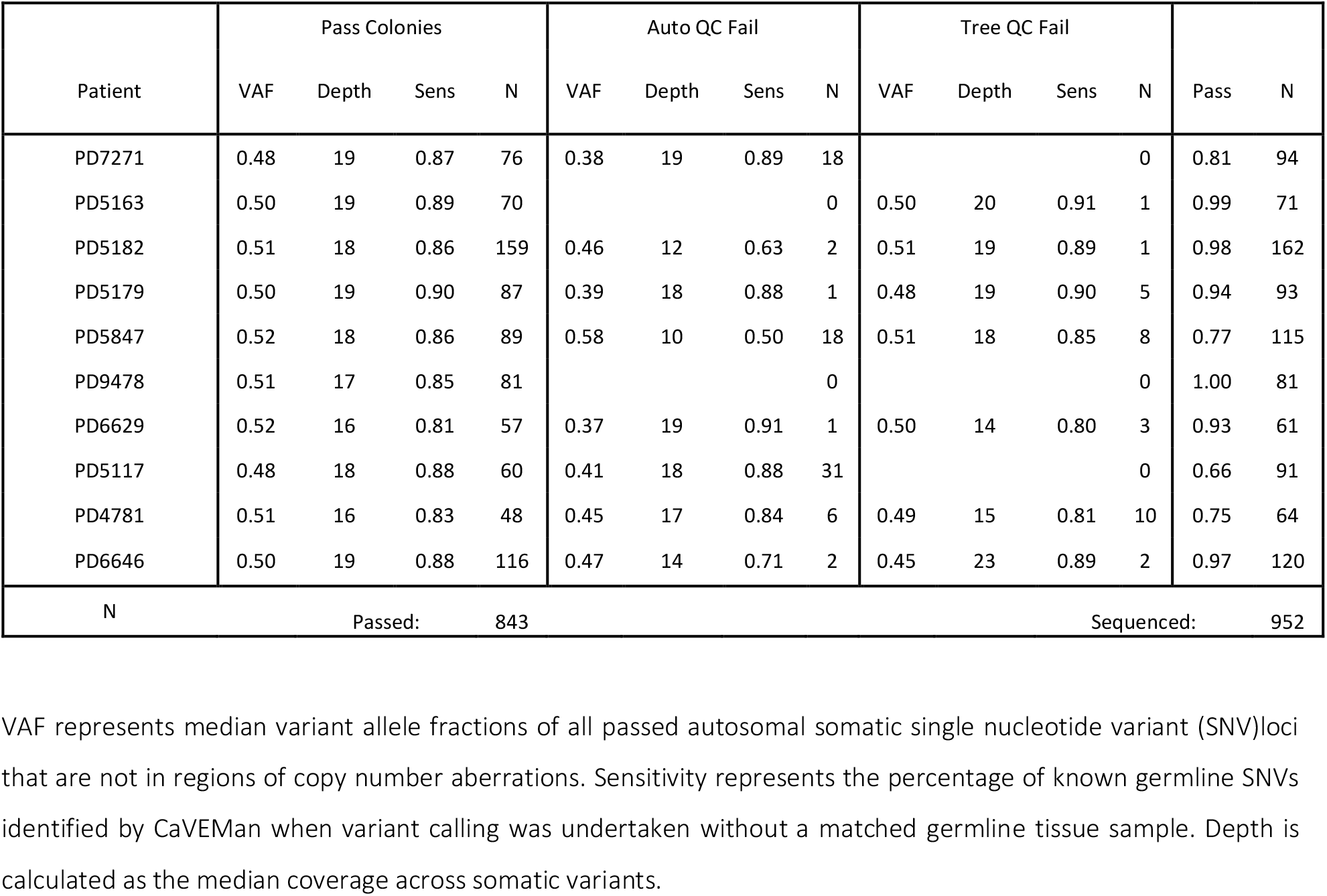
Number of colonies per patient sequenced and taken forward for analysis.

**Extended Table 3.**
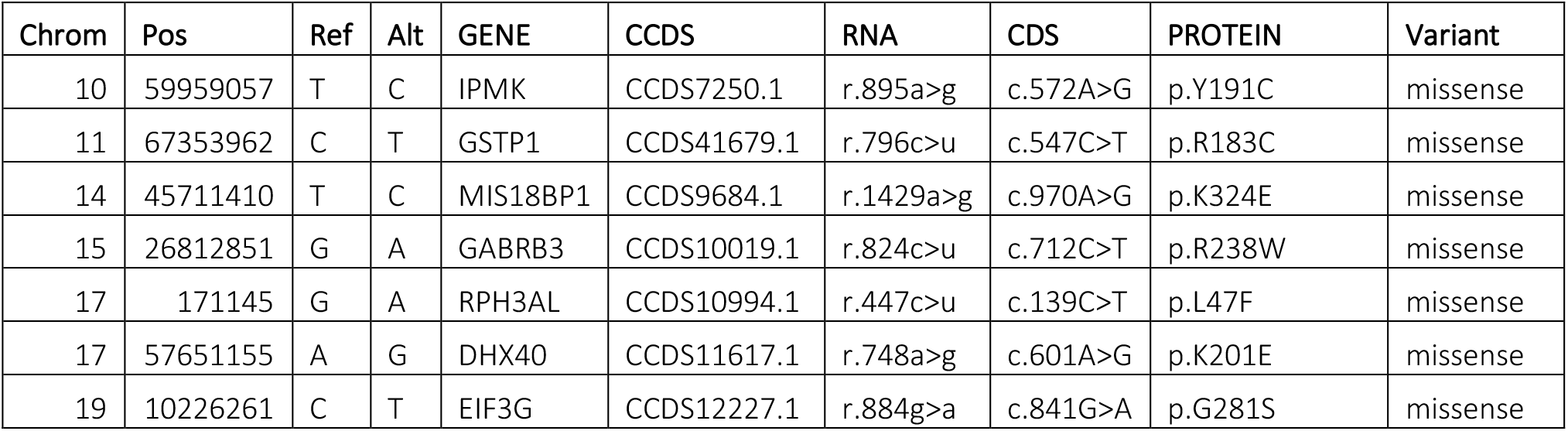
Coding mutations in the shared branch lacking a known driver mutation (PD6646).

**Extended Table 4.** Mutation burden and colony counts, including C>T at CpG sites across patients.

A.
| Subject | Driver Description | Number of SNVs | Colony Count |
| --- | --- | --- | --- |
| PD7271 | Wild Type | 27774 | 60 |
| PD7271 | JAK2 | 5664 | 16 |
| PD5182 | Wild Type | 82448 | 111 |
| PD5182 | JAK2/9pUPD + 1q+ | 10296 | 37 |
| PD5182 | 2pUPD | 593 | 1 |
| PD5182 | DNMT3A | 6743 | 10 |
| PD5182 | 9pUPD | 555 | 1 |
| PD5163 | Wild Type | 36752 | 55 |
| PD5163 | DNMT3A | 4352 | 9 |
| PD5163 | JAK2 | 3298 | 6 |
| PD5163 | chrX_Del | 649 | 1 |
| PD5847 | Wild Type | 11581 | 15 |
| PD5847 | JAK2/9pUPD | 16235 | 70 |
| PD5847 | DNMT3A | 3231 | 4 |
| PD5179 | Wild Type | 2808 | 3 |
| PD5179 | JAK2/+9 and 1q+ | 32205 | 84 |
| PD6629 | Wild Type | 896 | 1 |
| PD6629 | DNMT3A | 13808 | 19 |
| PD6629 | DNMT3A/JAK2 and TET2 | 13270 | 35 |
| PD6629 | DNMT3A | 1091 | 1 |
| PD6629 | CBL | 889 | 1 |
| PD9478 | Wild Type | 9548 | 8 |
| PD9478 | JAK2+DNMT3A | 23980 | 73 |
| PD5117 | Wild Type | 27982 | 22 |
| PD5117 | JAK2 | 36517 | 38 |
| PD6646 | Wild Type | 13510 | 17 |
| PD6646 | DNMT3A | 14722 | 22 |
| PD6646 | DNMT3A/JAK2 | 27252 | 76 |
| PD6646 | CBL | 655 | 1 |
| PD4781 | Wild Type | 1150 | 1 |
| PD4781 | JAK2 minor clade | 2101 | 2 |
| PD4781 | JAK2 major clade | 16129 | 45 |

B.
| Patient | Wild Type |  | JAK2 Branches |  | P<br>(JAK2>WT) | P_bonf | Phet | Phet_bonf |
| --- | --- | --- | --- | --- | --- | --- | --- | --- |
|  | C>T @<br>CPG | Not C>T @<br>CPG | C>T @<br>CPG | Not C>T @<br>CPG |  |  |  |  |
| PD7271 | 4571 | 23203 | 1021 | 4643 | 4.20E-03 | 4.20E-02 | 4.20E-03 | 4.20E-03 |
| PD5163 | 12992 | 69456 | 2336 | 7960 | 3.60E-71 | 3.60E-70 | 6.05E-68 | 6.05E-68 |
| PD5182 | 6481 | 30271 | 585 | 2713 | 9.00E-01 | 1.00E+00 | 2.10E-01 | 2.10E-01 |
| PD5179 | 1978 | 9603 | 3936 | 12299 | 6.83E-47 | 6.83E-46 | 2.26E-54 | 2.26E-54 |
| PD5847 | 400 | 2408 | 4223 | 27982 | 9.48E-02 | 9.48E-01 | 9.48E-02 | 9.48E-02 |
| PD9478 | 128 | 768 | 2377 | 10893 | 6.75E-03 | 6.75E-02 | 1.59E-14 | 1.59E-14 |
| PD6629 | 1507 | 8041 | 4668 | 19312 | 4.66E-15 | 4.66E-14 | 4.66E-15 | 4.66E-15 |
| PD5117 | 3789 | 24193 | 4820 | 31697 | 2.10E-01 | 1.00E+00 | 2.10E-01 | 2.10E-01 |
| PD4781 | 1994 | 11516 | 4067 | 23185 | 6.72E-01 | 1.00E+00 | 9.05E-01 | 9.05E-01 |
| PD6646 | 162 | 988 | 2332 | 15898 | 2.20E-01 | 1.00E+00 | 8.68E-06 | 8.68E-06 |

**Extended Table 5.** Rate of clonal expansion and timing of driver mutation acquisition.

| Patient | Clade | S<br>(Median) | 95% CI | dt<br>(Median) | dt CI.95% | N |
| --- | --- | --- | --- | --- | --- | --- |
| PD5847 | *JAK2*<sup>V617F</sup>, *9pUPD*, *TET2*<sup>N281fs*1</sup> | **2.33** | 1.43-3.60 | **33.03** | 29.14-35.44 | 937868 |
| PD5182 | *9pUPD* | **1.42** | 0.75-2.79 | **18.37** | 11.47-20.79 | 959453 |
| PD6646 | *DNMT3A*<sup>p?</sup>, *JAK2*<sup>V617F</sup> | **1.28** | 0.92-2.66 | **60.11** | 52.63-64.23 | 833501 |
| PD4781 | *JAK2*<sup>V617F</sup>, *9pUPD*, *TET2*<sup>Q1632*</sup>, *7q-*, *7p-* | **1.19** | 0.76-2.89 | **62.17** | 54.06-66.67 | 944472 |
| PD5179 | *JAK2*<sup>V617F</sup>, *1q+* | **1.15** | 0.86-1.70 | **27.99** | 23.32-30.52 | 928901 |
| PD6629 | *DNMT3A*<sup>R882H</sup>, *JAK2*<sup>V617F</sup>, *TET2*<sup>Q744fs*10</sup> | **0.83** | 0.36-1.66 | **43.83** | 37.29-48.71 | 910123 |
| PD9478 | *JAK2*<sup>Exon 12</sup>, *DNMT3A*<sup>Y908*</sup> | **0.71** | 0.54-0.96 | **42.90** | 35.72-47.82 | 884937 |
| PD6646 | *CBL*<sup>C401S</sup> | **0.70** | 0.34-3.69 | **48.65** | 35.65-55.00 | 726678 |
| PD7271 | *JAK2*<sup>V617F</sup> | **0.68** | 0.41-0.95 | **2.20** | 0.85-3.45 | 956233 |
| PD5847 | *JAK2*<sup>V617F</sup>, *9pUPD* | **0.67** | 0.06-2.46 | **22.80** | 5.91-27.40 | 921206 |
| PD5163 | *JAK2*<sup>V617F</sup> | **0.43** | 0.19-0.65 | **7.08** | 1.32-10.50 | 946952 |
| PD6629 | *DNMT3A*<sup>R882H</sup>, *JAK2*<sup>V617F</sup> | **0.40** | 0.27-0.56 | **24.68** | 15.70-29.47 | 811449 |
| PD5163 | *DNMT3A*<sup>T275fs*41</sup> | **0.38** | 0.22-0.61 | **5.85** | 1.03-9.66 | 941600 |
| PD6629 | *DNMT3A*<sup>R882H</sup> | **0.26** | 0.19-0.35 | **2.71** | 0.38-5.33 | 813866 |
| PD5117 | *JAK2*<sup>V617F</sup> | **0.18** | 0.13-0.23 | **8.74** | 2.13-13.35 | 860340 |
| PD5847 | *DNMT3A*<sup>Y660F</sup> | **0.09** | 0.05-0.25 | **0.23** | 0.12-0.95 | 838709 |

B.
| ACF (%) | Age (years) | Patient and clade |
| --- | --- | --- |
| 0.01 | 12 weeks (pc) | PD5847_DNMT3A |
| 0.1 | 3.87 | PD5847_DNMT3A |
| 1 | 29.70 | PD5847_DNMT3A |
| 10 | NA | PD5847_DNMT3A |
| 0.01 | 27.88 | PD5847_JAK2 |
| 0.1 | 32.86 | PD5847_JAK2 |
| 1 | 37.55 | PD5847_JAK2 |
| 10 | 42.37 | PD5847_JAK2 |
| 0.01 | 37.08 | PD5847_TET2 |
| 0.1 | 38.94 | PD5847_TET2 |
| 1 | 40.84 | PD5847_TET2 |
| 10 | 42.84 | PD5847_TET2 |
| 0.01 | 8.95 | PD7271_JAK2 |
| 0.1 | 13.30 | PD7271_JAK2 |
| 1 | 17.79 | PD7271_JAK2 |
| 10 | 22.29 | PD7271_JAK2 |
| 0.01 | 27.16 | PD5117_JAK2 |
| 0.1 | 41 | PD5117_JAK2 |
| 1 | 54.78 | PD5117_JAK2 |
| 10 | 68.86 | PD5117_JAK2 |
| 0.01 | 13.90 | PD6629_DNMT3A |
| 0.1 | 24.32 | PD6629_DNMT3A |
| 1 | 34.18 | PD6629_DNMT3A |
| 10 | 44.64 | PD6629_DNMT3A |

Rates of clonal expansion per year (selection, *S*) and the timing of driver mutation acquisition as inferred by approximate Bayesian computation. *S*, Selection (median); *dt*, median timing of driver mutation acquisition in post-conception years; *N*, number of simulations. Clones with sufficient immediate descendants (>5 branches) were selected for estimation of *S*. **B.** Trajectories of growth for the different mutant clades were estimated in patients and provided patient ages at which the mutant clones were at different cell fractions. Ages in years unless otherwise specified. ACF, aberrant cell fraction; pc, post conception.

